# Identifying triplex binding rules *in vitro* leads to creation of a new synthetic regulatory tool *in vivo*

**DOI:** 10.1101/2019.12.25.888362

**Authors:** Beate Kaufmann, Or Willinger, Noa Eden, Lisa Kermas, Leon Anavy, Oz Solomon, Orna Atar, Zohar Yakhini, Sarah Goldberg, Roee Amit

## Abstract

Nature provides a rich toolbox of dynamic nucleic acid structures that are widespread in cells and affect multiple biological processes^1^. Recently, non-canonical structures gained renewed scientific and biotechnological interest^2,3^. One particularly intriguing form of such structures are triplexes^4^ in which a single-stranded nucleic acid molecule interacts via Hoogsteen bonds with a DNA/RNA double helix^5^. Despite extensive research *in vitro*^6–9^, the underlying rules for triplex formation remain debated and evidence for triplexes *in vivo* is circumstantial^10–12^. Here, we demonstrate the development of a deep-sequencing platform termed Triplex-Seq to systematically refine the DNA triplex code and identify high affinity triplex forming oligo (TFO) variants. We identified a preference for short G-rich motifs using an oligo-library with a mix of all four bases. These high-information content motifs formed specific high-affinity triplexes in a pH-independent manner and stability was increased with G-rich double-stranded molecules. We then conjugated one high-affinity and one low-affinity variant to a VP48 peptide and studied these synthetic biomolecules in mammalian cells. Using these peptide-oligo constructs (POCs), we demonstrated possible triplex-induced down-regulation activity in 544 differentially expressed genes. Our results show that deep-sequencing platforms can substantially expand our understanding of triplex binding rules, which in turn has led to the development of a functional non-genetically encoded regulatory tool for *in vivo* applications.

---

Shortly after Francis Crick and James Watson published the iconic DNA double-helix model, other nucleic acid structures were detected. Most notably the existence of triple helical structures (triplexes) was first supported in a 1957 study^4^. The underlying interactions between a third strand and the double helix are based on Watson-Crick-independent hydrogen interactions and are termed Hoogsteen bonds^5^. In triplexes, single-stranded chains bind the polypurine stretch of the major groove of the duplex molecule (dsDNA) via two possible Hoogsteen configurations: (i) Hoogsteen interactions promoting a parallel orientation (pyrimidine motif) and (ii) reverse Hoogsteen bonds resulting in an anti-parallel orientation (purine motif) of the third strand with respect to the duplex sequence. Given the specificity in formation of triplexes^6,7,9^, Moser and Dervan^8^ developed short DNA-based triplex-forming oligonucleotides (TFOs) that are typically 15-30 nt long and form triplexes with polypurine stretches of the dsDNA termed triplex target site (TTS). Due to their specificity in targeting dsDNA, TFOs were subsequently used to identify triplex rules^13,14^ and applied as biotechnological tools *in vitro* and *in vivo* to regulate transcription^15–18^, control recombination events^19^ and induce triplex-directed mutagenesis^20–22^. Despite extensive amount of work to establish TFOs as therapeutic agents, triplex formation *in vivo* mostly was attenuated and the expected biological function was limited^23,24^.

To date, there is an insufficient understanding of the underlying ‘triplex code’. It is unclear what the minimum length of nucleotide stretches needed to from triplexes is, how purine/pyrimidine mixes impact triplex formation, and how many mismatches can be tolerated within a triplex-forming sequence. This problem is further compounded by having only a few examples for TTS sequences that have been verified to form triplexes *in vitro*. As a result, the need to match a 15-30 nt sequence specifically creates a large search space which is difficult to explore with traditional methods. This has led to a sub-optimal understanding of what motifs are required to form high-affinity TFO-TTS pairs, which in turn has limited our ability to either find evidence for the existence of triplex formation *in vivo* or make use of synthetic TFOs for various i*n vivo* applications. In addition, triplex formation has implications for various other research fields including material sciences through developing triplex-responsive hydrogels^25^, synthetic biology-inspired diagnostics^26^ and biosensors^27^, nanotechnology-based switches^28,29^, ideas to expand the DNA origami toolbox^30^, and DNA storage. Finally, with the growing interest in dynamic, non-canonical structures, and their hypothesized relation to diseases^31^ such as the ones associated with nuclear repeat disorders (e.g. Fragile X, Friedreich Ataxia, Huntington’s disease, etc.), a detailed understanding of the underlying code of triplex formation must be at the forefront of these endeavors, and is thus urgently needed.

In this work, we employ a high-throughput oligo library and next generation sequencing based approach to substantially expand the space of known TFO partners to a set of G-rich TTS sequences, thus allowing us to extract new binding rules in the form of triplex forming motifs (TFMs). We then selected two variants (high and low-affinity) and studied them in an *in vivo* setting using either a native oligo form or as a conjugate to a regulatory peptide. Our results show that our high affinity TFO variant in its conjugated form was able to massively disrupt expression of hundreds of genes likely as a result of triplex interaction with the genome, thus demonstrating a new TFO-based tool for regulatory applications.

## Results

### Triplex-seq

We developed a technology platform based on DNA synthesis and next-generation sequencing termed Triplex-Seq to study triplex formation *in vitro*. To do this, we combined an electrophoretic mobility shift assay^32^ (Figure 1a) with Illumina sequencing and DNA synthesis technologies (Figure 1b). Briefly, the Triplex-Seq platform comprises a (i) single variant of a triplex-target site (TTS) and (ii) library of mixed-based triplex-forming oligos (Figure 1c). Triplex formation between the TTS and the TFO library is induced in parallel (henceforth referred to as pH 5) or anti-parallel conditions (henceforth referred to as pH 7) and the products (triplex, TTS, TFO) are separated on a native polyacrylamide gel (PAGE). The TFO variants that were mixed with the TTS (triplex lane) or without TTS (TFO-only lane, background) are extracted from a position on the gel which coincides with the shifted triplex band, the triplex is disrupted and TTS discarded and the TFOs from both lanes are sequenced. To screen for single-stranded TFO sequences that formed triplexes, we designed TFO libraries (Figure 1c) using the randomized or mixed-base tool from integrated DNA technologies (IDT). Mixed bases are represented by the IUPAC (International Union of Pure and Applied Chemistry) single-letter code. Using the standard machine mixing, degenerate bases are incorporated into the TFO sequences at the wobble/mixed-base positions (e.g. for the N-TFO library, guanine (G)/adenine (A)/thymine (T)/cytosine (C) are incorporated with a 1/1/1/1 ratio) creating a diverse TFO library. Using this randomized TFO library generation, 13 small (27 variants) to large (270,000,000 variants) TFO libraries were designed (Figure 1d).

**Figure 1.**
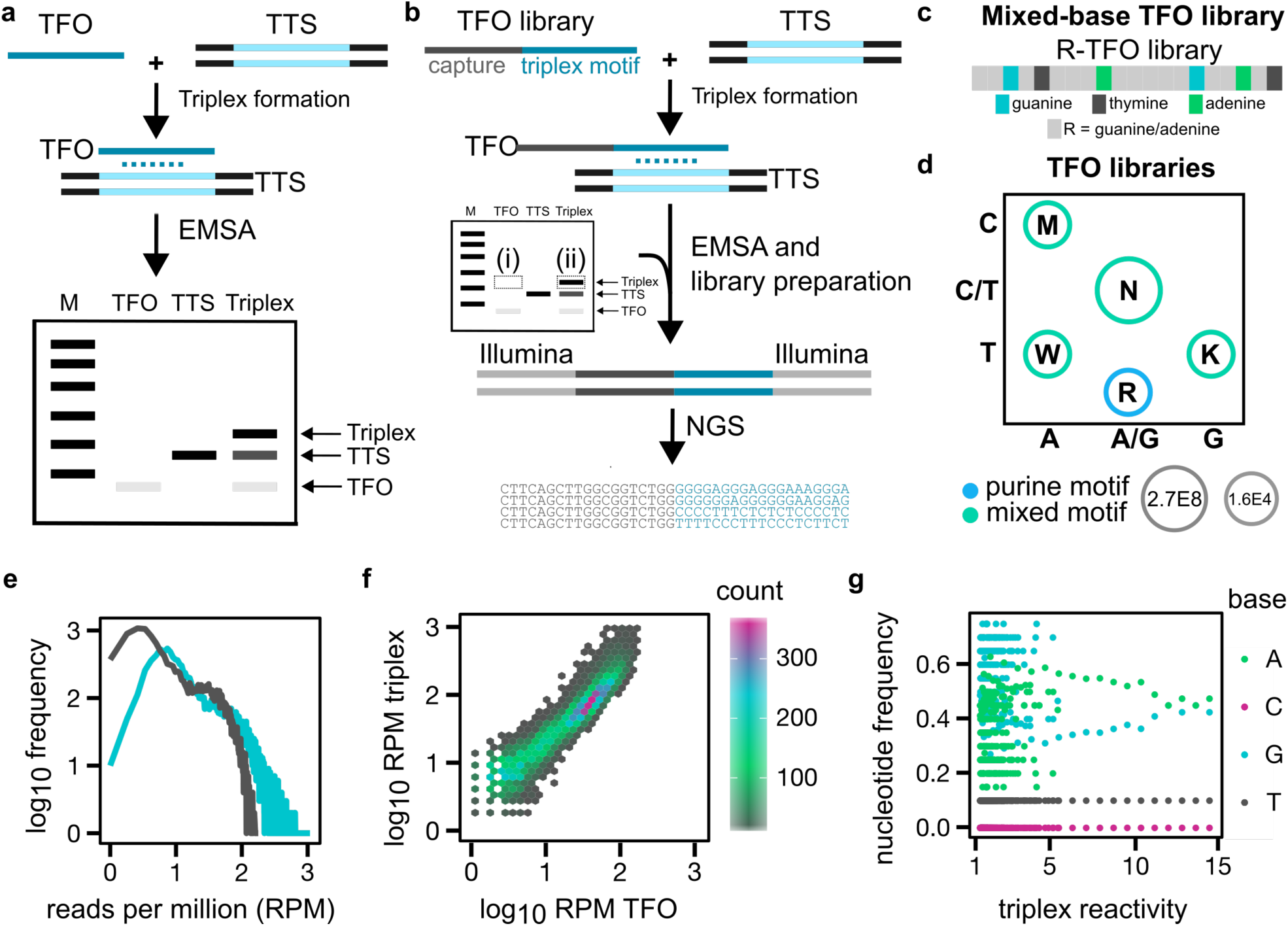
Schematic representation of Triplex-Seq design and analysis. **(a)** An electrophoretic mobility shift assay (EMSA) is classically used for triplex formation experiments. Single-stranded triplex forming oligos (TFOs) are mixed with a double-stranded triplex target site (TTS) in triplex-favoring conditions. The products are separated on a native 10-20 % polyacrylamide gel (PAGE) and migration of TFO, TTS and triplex is visualized. A shift between the faster-migrating duplex and slower-migrating triplex is expected, as schematically shown. **(b)** In the Triplex-Seq platform, variants of a TTS are mixed with a library of mixed-base TFOs. The TTS (between 30-80 bp long) harbors the purine-rich segment that can accommodate a third strand. The short TFOs (up to 30 nt) contain the putative DNA stretches that form triplexes with the TTS in a parallel or anti-parallel orientation. After incubation, the products were separated on a 10 % PAGE and bands corresponding to the triplex position were cut from the gel. Extracted TFO sequences were prepared for next-generation sequencing (NGS) via PCR amplification and subsequently bioinformatically analyzed. **(c)** The mixed-base TFO libraries consist of a 19 nt long capture sequence serving as a PCR primer docking site common to all of the TFO libraries, 6-7 fixed bases (either G, A, T or C) that serve as an internal barcode, to identify the TFO sequences in the sequence reads as well as 13-14 mixed bases which are incorporated at various ratios. We show the R-TFO library as an example. (**d)** A variety of TFO libraries were generated and are shown as a function of nucleotide content and library size. **(e)** Example analysis of R-TFO in which the frequency of normalized read counts is plotted against the RPMs for both TFO-only (grey) and triplex lanes (blue). (**f)** Hexagonal scatter plot depicting RPM values for TFO-only and triplex lanes. Bins are color-coded by RPM counts. **(g)** Plot depicting nucleotide frequency within TFO as a function of triplex reactivity score. Note the increase in G percentage as a function of reactivity score. See methods for TTS used in this experiment. All experiments were carried out in duplicates on separate days.

In Figure 1e-g we plot the analysis pipeline for the example R-TFO library (G/A) in pH 7. To compare datasets, we first computed the normalized read counts (reads per million, RPM) of the TFO and triplex lane, and plotted the frequency of RPMs (Figure 1e). We observe a decay for the TFO lane in which most TFO sequences that were identified appear once or twice (grey line), while in the triplex lane a subset of reads appears with higher frequency (blue line) indicating enrichment. In Figure 1f, we plot the RPMs of the TFO lane vs the RPMs of the triplex per variant. Here, we divide groups of variants into bins defined by the relevant range of RPMs per variant for the TFOs and triplexes. All bins above approximately 2.3 represent sequences whose RPM value is larger than the maximal RPM value obtained for the TFO-only lane. These are, therefore, highly enriched variants. To further characterize the sequences, we introduce a new score termed ‘triplex reactivity’ which is defined as: 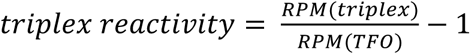. A positive reactivity score indicates an enrichment of a particular variant above the TFO-only control. In Figure 1g, the mean nucleotide frequency of each nucleotide (G/A/T/C) within the TFO sequences at a given triplex reactivity value were plotted as a function of the triplex reactivity scores. For the example R-TFO library, we observe that cytosines and thymines are absent or stay constant throughout each triplex reactivity score while adenines decrease and guanines increase with increasing triplex reactivities.

### Triplex-seq reveals new triplex forming motifs (TFMs)

We first tested TFO libraries that contained a mixture of only two bases in each of the 14 variable positions (K, M, R, W) in pH 5 and pH 7 (Figure 2a). The distribution and probability density of all triplex reactivity values for both buffer conditions and for each of the TFO libraries is shown in **Error! Reference source not found.**b and Extended Data Figure 1a. A higher frequency of triplex reactivity values (weak triplex formation) ranging at around 0 is observed in pH 5 except for the M-TFO library (A/C), whereas triplex reactivities for all TFO libraries tested in pH 7 are higher indicating that the pH 7 conditions are more permissive for triplex formation. Next, to search for enriched TFMs that can be found in the most reactive TFO sequences, we used DRIMust^33^, a tool that identifies enriched k-mers and motifs based on a ranked list of sequences. Here, we applied DRIMust on sorted triplex reactivity lists to detect k-mers and enriched consensus TFMs that are significantly over-represented at variants with high-reactivity scores. DRIMust was applied to every one of the four TFO libraries and the consensus TFMs are shown in Figure 2c. We note that all TFMs range between 5-10 nt. For pH 5, a consensus TFM was only detected for the M-TFO library, whereas consensus TFMs for all TFO libraries were found for pH 7. In cases where no motif was identified, the algorithm was not able to detect enriched k-mers in the sorted list at a minimum hypergeometric (mHG) p-value threshold of 10^−6^. By comparing our enriched TFMs to past literature findings, we obtain support for the sensitivity and validity of the Triplex-Seq platform. Specifically, the R-TFO library (G/A) displays a short GARA-TFM (p-value ≤ 7.7×10^−24^). which has been associated in the Friedreich’s Ataxia disease^34^ and was found to downregulate expression of the *dhfr* gene^35,36^. The K-TFO library (G/T) features a short consensus TFM with 80 % thymines (p-value ≤ 8.7×10^−27^). Formation of triplexes with mixed GT-TFMs have been described to form stable triplexes in acidic and neutral pH^37,38^. The W-TFO library (A/T) exhibits stretches of adenines as well as thymines (p-value ≤ 4.9×10^−324^). Both nucleotides have been shown to form triplets with an A-T basepair while adenines also interact with a G-C basepair^37^. The M-TFO library (adenine/cytosine) reveals an A/C mixed-consensus TFM (pH 5, p-value ≤ 1.4×10^−134^; pH 7, p-value ≤ 7.2×10^−230^) in both conditions. When looking at the top hits of the ranked triplex reactivity lists (Supplementary Table 1), a preference for adenines can be observed (60 – 70 %) which can be explained by adenine’s ability to bind to both G-C and A-T basepairs. The A/C-mixed TFMs however might result from formation of non-canonical triplexes by binding to the C-T and T-A basepairs as was proposed previously^37^ or via the potential formation of parallel triplexes due to cytosine protonation at neutral pH^39,40^. To further support this finding, triplex formation was confirmed by EMSA using an A-rich variant from M library at pH 5 conditions with one of the highest triplex reactivity scores (Extended Data Figure 1b).

**Figure 2.**
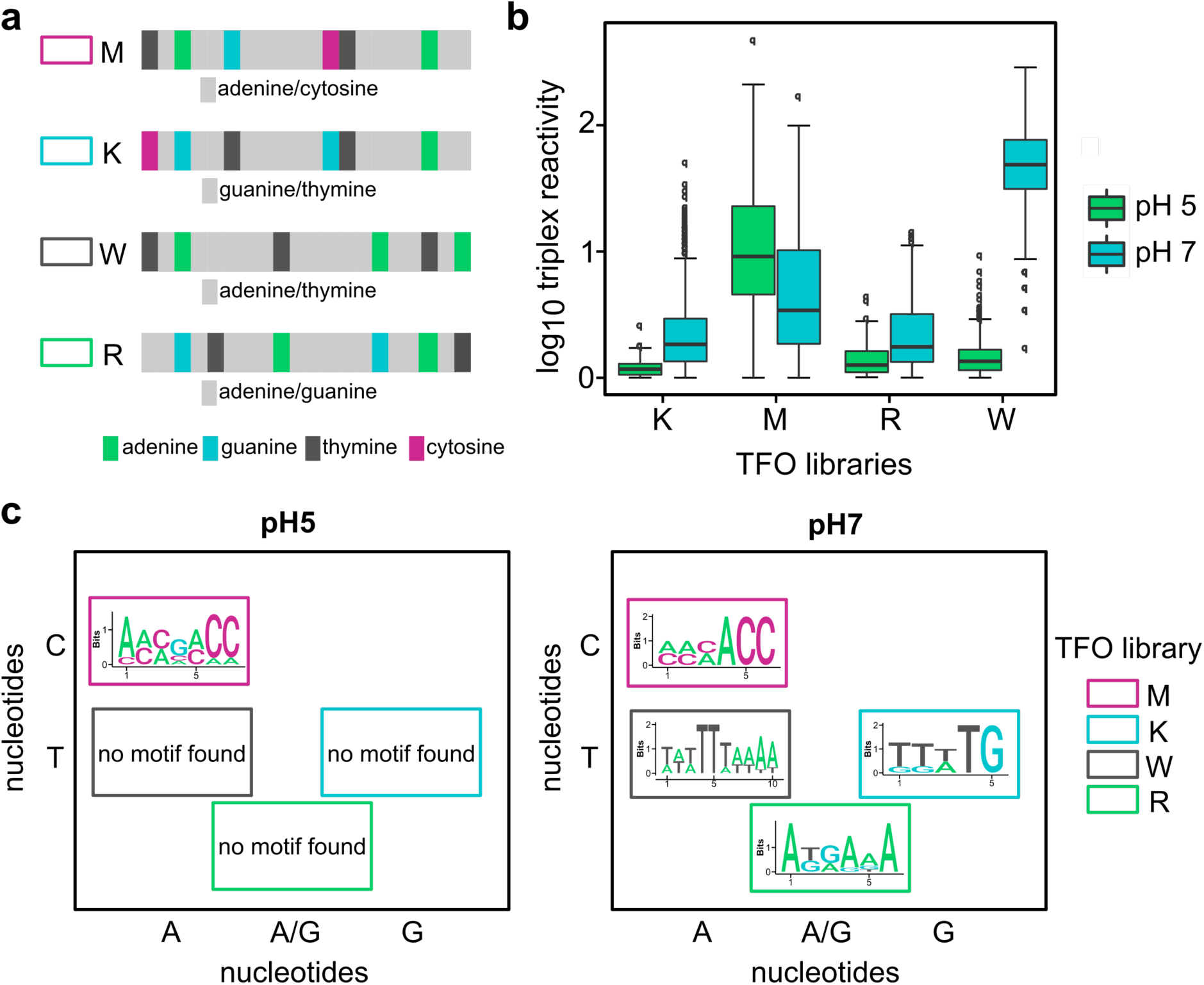
pH-dependent triplex formation using small TFO libraries. **a**, The enrichment of sequences of TFO libraries with 2-mixed bases (K, M, R, W) were tested using the Triplex-Seq platform. Each library contains 1.6×10^4^ variants and was tested in pH 5 and pH 7, respectively. **b**, Distributions of triplex reactivities of the four TFO libraries that were tested in pH 5 (green) and pH 7 (blue). **c**, Analysis of sequence preferences was carried out using DRIMust^33^. DRIMust k-mers (Supplementary Table 1) and motifs were computed based on ranked triplex reactivity lists. While consensus motifs of 5-10 nt were detected for all libraries in pH 7 (with high triplex reactivity values), a DRIMust consensus motif was computed only for the M-TFO library in pH 5.

### G-rich TFOs are enriched in a random TFO library

We next sought to evaluate which TFO sequences will be enriched when testing the ∼ 270,000,000 large N-TFO library (Figure 3a, top). We used two known TTSs (pH 5: TTS 2)^41^ and (pH 7: TTS 1)^24^ as our test targets (Figure 3a, middle and bottom) alongside the N-TFO library to determine the migration pattern of duplexes and triplexes in various buffer conditions. In Figure 3b, we plot the mean nucleotide frequency of the enriched TFO sequences against the triplex reactivities for pH 5 (Figure 3b, left) and pH 7 (Figure 3b, right) and observe a G-increase in the TFO sequences with higher triplex reactivities for both conditions. To further characterize the behavior of the N-TFO, we additionally applied the Triplex-Seq platform in a buffer that negatively affects triplex formation due to the presence of monovalent ions (Figure 3c)^24,42^. In Figure 3d the triplex reactivities of both TTS sequences in the triplex-disfavoring buffer are plotted and we observe again a trend toward G-rich sequences with higher triplex reactivities, but this effect is weaker as compared to the conditions lacking potassium (Figure 3b). While it is commonly assumed that physiological concentrations of potassium abolish triplex formation, more sensitive assays showed that triplex formation is reduced to 10-20 % triplexes compared to triplex-favoring conditions^23,43,44^. Comparing the triplex reactivities of all conditions shown in Figure 3e (left) a similar decrease of the triplex reactivity values was observed when comparing the mean triplex reactivities from pH 7 with pH 5 (46 %) and pH 7+K^+^ (TTS 2: 62 %, TTS 1: 56 %) suggesting that more stable triplexes are formed in neutral compared to acidic or neutral pH with high potassium concentrations. To confirm that TFOs with high triplex reactivities form triplexes, single variants with high triplex reactivities were tested for their triplex formation potential EMSA (Extended Data Figure 2a) as well as circular dichroism (CD) spectroscopy (Extended Data Figure 2b-d). Both assays confirmed that stable triplexes form with TFOs exhibiting high triplex reactivities.

**Figure 3.**
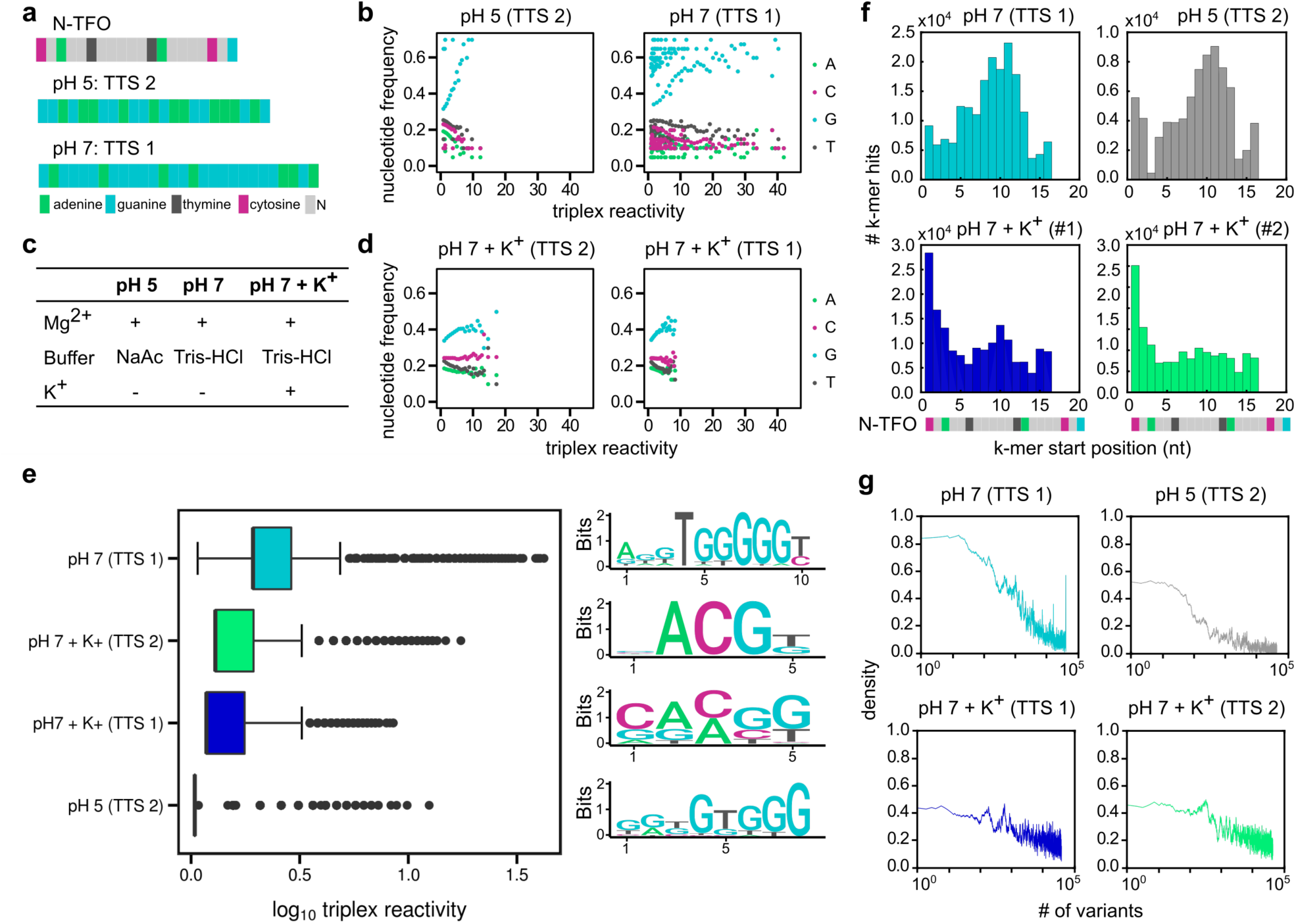
Identification of G-rich TFO motifs using N-TFO library. **a**, A 270,000,000 variant large N-TFO library was tested with two single variants of triplex target sites (TTS 1 in pH 7, TTS 2 in pH 5) in triplex-favoring conditions (pH 5 and pH 7). **b**, Mean nucleotide frequency as a function of triplex reactivity values in triplex-favoring buffer conditions are plotted for pH 5 (left) and pH 7 (right). In both plots green, magenta, gray, and light blue circles correspond to adenine, cytosine, thymine, and guanine frequencies, respectively. **c**, Table containing triplex favoring (left and middle column) and disfavoring (right column) conditions **d**, Mean nucleotide frequency as a function of triplex reactivity values in triplex-disfavoring buffer conditions are plotted for pH 5 (left) and pH 7 (right). **e**, (Left) comparison of triplex reactivity distributions for all four buffer-TTS combinations: pH7-TTS1 (light blue), pH7-TTS2 triplex disfavoring (green), pH7 – TTS1 triplex disfavoring (blue), pH5-TTS2 (gray). (Right) Associated logos for an enriched motif extracted via DRIMust for reach respective reactivity distribution plotted on the box. **f**, In the DRIMust k-mer-position analysis, it can be observed that for the triplex-favoring conditions (pH 5 and pH 7), the enriched k-mers cluster in the center of the TFO sequence, whereas the position of the k-mers of the triplex-disfavoring are enriched in the beginning of the TFO sequence. **g**, k-mer density plotted as function of the number of variants from the sorted triplex reactivity list for TTS1-pH 7 (left-turquoise), TTS2-pH 5 (left-gray, TTS1- triplex disfavoring (right-blue), and TTS2-triplex disfavoring (right-green).

Applying DRIMust to the N-library reactivity sorted lists reveals consensus TFMs that display a clear pattern of G-stretches within a 7-10 nt long consensus motif for acidic (p-value ≤ 10^−82^) and neutral pH (p-value ≤ 2×10^−298^) without potassium (Figure 3e, bottom and top logos, respectively). However, in contrast to this finding a 5 nt long ACGT TFM for both TTS sequences in the potassium-containing conditions is observed (Figure 3e, middle logos). This suggests that in pH 7+K^+^ conditions a mix of non-canonical structures of weakly interacting TFOs (i.e. forming unstable triplexes) and G-rich TFOs that form G-quadruplexes migrate through the gel, which together result in the ACGT-consensus motif (p-values ≤ 2×10^−53^). This is consistent with previous findings where stable G-rich triplexes have been described *in vitro* ^23,32,45,46^ and *in vivo* ^20,47^, but were also found to form G-quadruplexes in physiological environments^2,32^. Furthermore, comparison of the positions of the enriched k-mers within the TFOs for the triplex-favoring and triplex-disfavoring conditions showed that the G-rich k-mers for the pH 5 and pH 7 samples cluster in the center of the TFO sequence, while a preference for the beginning of the TFO sequence was observed for the triplex-disfavoring condition (Figure 3f). To further study the significance of the DRIMust TFMs, we examined the density of the enriched k-mers within the sorted list (see Supplementary Table 1) as function of variant position in the sorted triplex reactivity lists by computing a running average of the number of k-mers in a 100 variant window (Figure 3g). The data shows that for the triplex favoring conditions (two left panels) the density exhibits a step-like response with a relatively high values for a small subset of variants at the top of the list, which rapidly drops off to a small value for the remaining variants. Conversely for the triplex disfavoring conditions a gradual decline in density is observed as a function of variant position in the list. Thus, the position distribution of k-mers within the TFOs together with the k-mer density plots support an interpretation that TFMs that are associated with triplex formation exhibit three distinct signatures that allow us to separate them from triplex disfavoring controls: higher p-values as compared with triplex disfavoring controls, a distinct step-like density function, and logo position distributions that are centered in the middle of the TFO sequences.

### G-rich TFOs form stronger triplexes with G-rich TTSs

We next studied the relationship between TFO enrichment and variation in guanine/adenine ratios in the TTS for triplex formation. Five single TTS variants were designed *de novo* in which the ratio of guanine to adenine was systematically changed (20/80 to 80/20) (Figure 4a and Extended Data Figure 3) and tested with the N-TFO library in pH 5 and pH 7 (Figure 4a, top). We observe the expected exponential decay for the TFO-only lane and an increase in the number of enriched TFO variants for the triplex lanes (Figure 4b) as well as a slight increase in triplex reactivities (Figure 4c), that correlated with a higher percentage (75 – 80 %) of guanine within the TTS, supporting the importance of G-rich TTS sequences for stable triplex formation in both pH 5 and pH 7. To evaluate the enriched TFO sequences, the mean nucleotide frequency of the four bases was computed and is plotted in Figure 4d. In the heatmap, each line corresponds to one TTS variant and each column represents the triplex reactivity that was divided into three ranges (low, medium, high). The heatmap displays the mean nucleotide frequency of TFO sequences in the respective triplex reactivity range. The data shows that high-reactivity TFO variants are characterized by increasingly G-rich TFO sequences. For pH 7, this effect becomes more pronounced with increasing percentage of guanines in the TTS. This increase in G-content for the TFO comes at the expense of all other nucleotides, and is not restricted to a reduction in the presence of a particular nucleotide.

**Figure 4.**
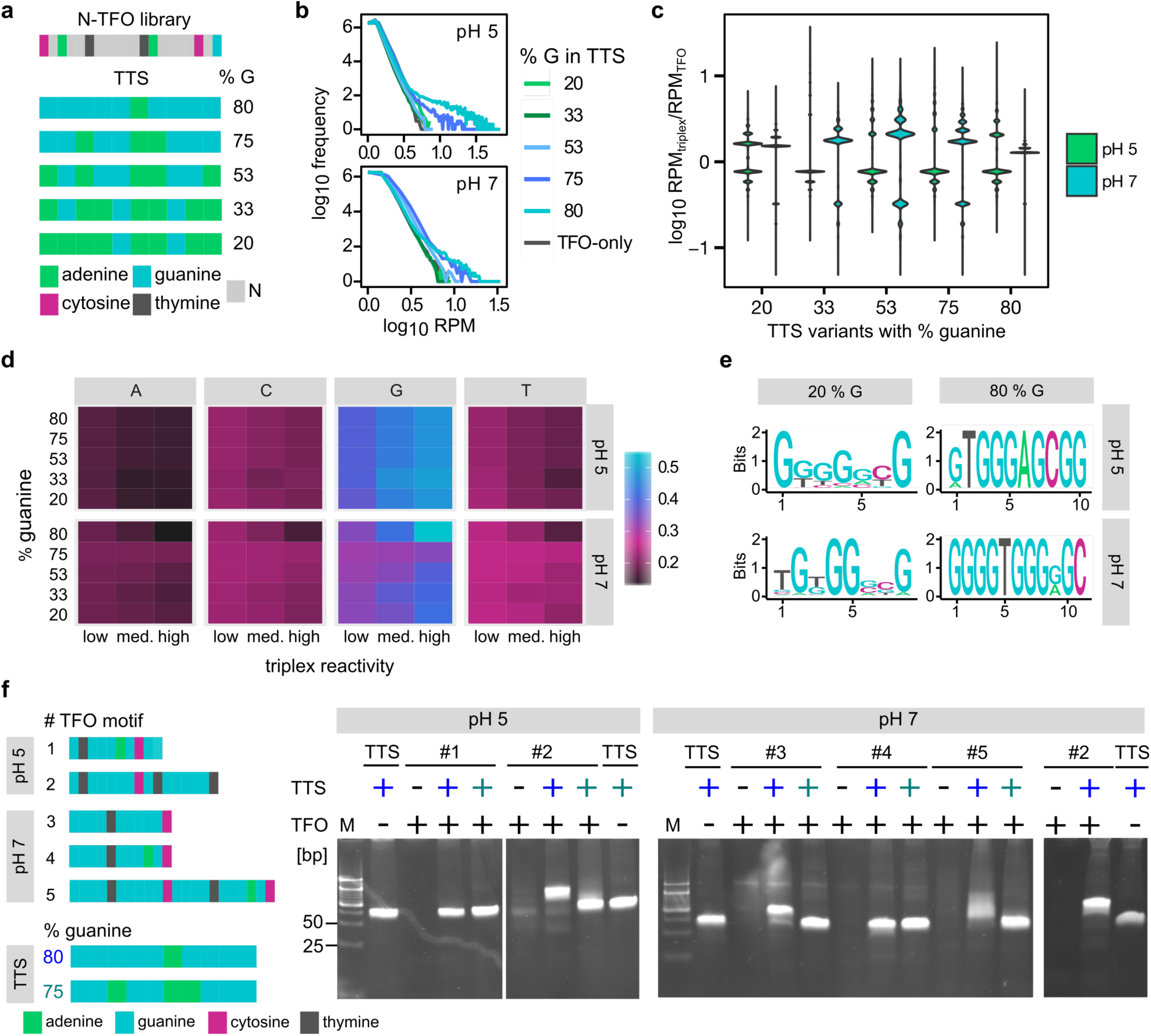
High guanine content positively influences triplex formation. **a**, Schematic showing the TFO (top) and the five TTSs used in the experiments below. **b**, Frequency of the RPM counts is plotted against the RPMs for each TTS (20%G (light green), 33%G (dark green), 53%G (light blue), 75%G (blue), 80% (turquoise)) for pH 5 (top) and pH 7 (bottom) conditions. **c**, Violin plot of the distributions of the ratio of RPM_triplex_ and RPM_TFO_ for all five TTSs at pH 5 (green) and pH 7 (light blue) conditions. **d**, Heatmap depicting the nucleotide frequency as a function of triplex reactivity and experimental conditions: pH 5 (top) and pH 7 (bottom). Each column represents a range of triplex reactivity values (low: < 1.7, medium: 1.7-2.6, high: >2.5), and each line corresponds to a TTS variant. The nucleotide frequency of each nucleotide is colored (right bar). **e**, Sample DRIMust consensus motifs for: pH 5 and TTS20 (top-left); pH 5 and TTS80 (top-right); pH 7 and TTS20 (bottom-left); and, pH 7 and TTS80 (bottom-right). **f**, Triplex verification experiments. (Left - top) schematic for single the TFOs used in the EMSA experiment, and the associated TTSs (left-bottom). (Right) gel shift experiments showing putative triplex shifts for variants #2 (pH5 and pH7), #3, and #5.

To further strengthen this conclusion, we applied DRIMust to compute the enriched TFM logos that are associated with each TTS (Figure 4e and Extended Data Figure 4). The data shows that all the logos obtained display a G-rich TFM. However, with higher G content in the TTS, the logos also display an increasing amount of information. Specifically, TTS variants containing 20-53 % guanine in both pH 5 and pH 7 generate enriched TFMs containing two to three fixed bases (2 bits of information), which flank a stretch of five variable bases in the center of the logo containing less than 1 bit of information per position (Extended Data Figure 4). Conversely, for TTS variants with high guanine percentage (75-80 %, Figure 4e, right and Extended Data Figure 4), the TFM logos display fixed letters (2 bits of information) at nearly every position revealing a much higher information content in these motifs. The TFMs obtained for the 80 % TTS for both pH 5 and pH 7 conditions represents a strong preference for only a few TFO variants that are similar in sequence. This suggests that these TFO variants interact with the TTS molecules in both a reactive and extremely specific fashion. To confirm this prediction, we carried out a set of single variant EMSA experiments (Figure 4f). We used five TFOs and two TTSs (75% and 80% Guanine – Figure 4f -left-bottom). The image shows that we observe a triplex-shift for only the 80% Guanine TTS, which differs from 75% Guanine TTS by only two bases, and for 3 of the 5 TFO variants. In particular, a single mutation from guanine to adenine in the 8^th^ position (TFO #3 and #4) was sufficient for abolishing triplex formation. Therefore, triplex formation with the 80% Guanine TTS seems to only occur with a limited set of TFOs, especially if a single cytosine was present in the sequence.

Finally, we examined the density of k-mers identified in the sorted triplex reactivity lists (Extended Data Figure 5). The plots show a sharp step function when using G-rich TTS variants consistent with a stable and specific triplex interaction between the TTS and the TFOs. However, for TTSs with lower guanine concentration a gradual decline is observed similar to what was observed for the triplex disfavoring conditions (Figure 3f), which is consistent with an unstable and weak triplex interaction. These observations are in accordance with findings from another study which showed that G-rich TTSs (with ∼80% G-content) are necessary for triplex formation^13^. Hence, our results suggest that G-rich TTS variants lead to a stronger, more specific and pH-independent triplex interaction with TFO libraries that contain a few highly-specific variants, compared to TTS variants with lower guanine content (Figure 4f, lane 8, pH 5) which form weaker triplexes. Finally, the 7-10 nt long DRIMust G-rich consensus motifs identified in acidic and neutral pH prompted us to design TFO libraries (B- and D-TFOs) with a continuous stretch (3-9 nt) of mixed bases embedded in a constant non-G-rich background for the purpose of characterizing the minimal sequence length required for triplex formation (Extended Data Figure 6). Our observations show that only for the libraries with a 9 nt variable segment is a G-rich logo extracted, which suggests that a length of 9 nt is sufficient for triplex formation while shorter TFO sequences likely lack a strong triplex-formation potential.

### TFOs exhibit evidence for triplex functionality in CHO cells

In order to test whether the hi-affinity TFOs that were enriched *in vitro* can regulate expression in a biological setting, we selected two TFO variants. The first is a high triplex reactivity variant (pTFO, variant #3 in Extended Data Fig. 2) containing a high information content TFM, while the second was characterized with a low (∼5) reactivity score (nTFO) in Fig. 3b (top-right). Since activation and repression of promoters in eukaryotes are typically mediated by protein domains, we also ordered the same TFOs as peptide-oligo conjugates (POCs, from Bio-Synthesis, Inc.), with the TFO conjugated to three repeats of the transactivation domain of virion protein VP16, termed VP48 (Figure 5A). We additionally ordered an AlexaFluor488 fluorophore on the nTFO-VP48 (nPOC) and the nTFO with FAM modification (oFAM, from IDT) in order to quantify transfection efficiencies of TFOs and POCs.

**Figure 5.**
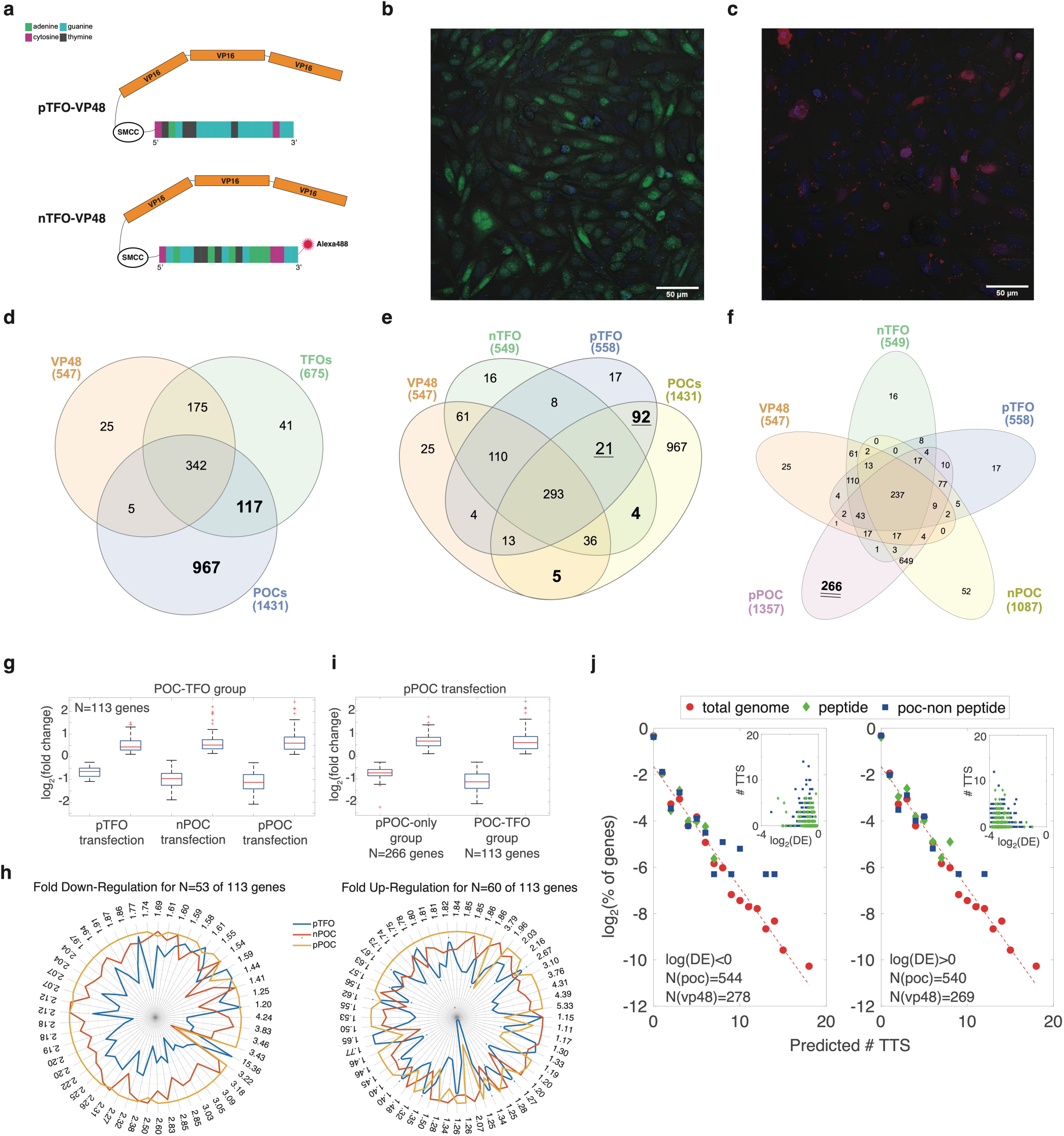
In vivo regulatory function exhibited by a high-reactivity TFO. (a) The two peptide-oligo conjugates used in the experiment. For both conjugates a peptide consisting of three repeats of VP16 was conjugated via the SMCC modification to the 5’ end to the TFO. (Top) pTFO-VP48 (pPOC) consisting of the verified and high triplex-reactivity #3 TFO in Figure 4f. (Bottom) nTFO-VP48 (nPOC) consisting of a low triplex reactivity TFO found in our in vitro experiments and an additional Alexa488 fluorescent label on the 3’ end. (b-c) Epi-fluorescent images of CHO/K1 cells transfected with nPOC (b) and oFAM (c) showing fluorescent nuclei 24 hours post-transfection. Scale bar corresponds to 5 μm. (d-f) Venn diagrams comparing the differentially expressed genes in different sub-group types: (d) Three sub-groups: peptide, TFO, POC. (e) 4 sub-groups: peptide, nTFO, pTFO, and POCs. (f) All 5 groups corresponding to the different transfection experiments. Note the especially high number of unique differentially expressed genes for pPOC. (g) Box plot depicting a comparison of log2(fold change) distributions for the 113 genes making up the POC-TFO group between the the pTFO, nPOC, and pPOC transfection experiments. (h) Radar plots showing the fold down- regulation (left) and up-regulation (right) for the genes in the group of 113. In each plot, each axis corresponds to a gene. The values on each axis ranges from 1 to the number written on the edge of the axis. (i) comparison of the POC-TFO and pPOC-only groups within the pPOC transfection experiment. (j) Predicted # TTS distribution exhibiting an exponential behavior. Genomic distribution and fit (red) overlaid by the distribution computed for the genes down-regulated (left) and up-regulated (right) in the POC experiment (blue squares) and not in the peptide control (green diamonds). (Inset) down (left) and up (right) regulated genes differentially expressed for both the peptide and POC groups are plotted as a function of predicted # of TTS.

We transfected Chinese hamster ovary cell line K1 (CHO/K1) (see Methods) with equal molar amounts of either nPOC or oFAM. After 24 hours, we performed Hoechst-staining and imaged the nPOC and oFAM transfected cells, as well as a non-transfected control, using a confocal microscope (Fig. 5b-c) (see Methods). In both images the Hoechst staining is demarked in blue, while the AlexaFluor488 labelled nPOC is shown in green (Fig. 5b), and the oFAM is depicted in red (Fig. 5c). The images show that 24 hrs after transfection, the nPOC accumulates within cell nuclei whereas the oFAM is barely visible. This indicates that the conjugation of the peptide to the 5’end of the oligo has introduced a stabilization effect by allowing the oligo (labelled by the fluorophore at the 3’end) to be stably present within the cell nuclei for an extended duration of time.

We further tested the various TFOs, POCs, as well as the VP48 peptide (Bio-Synthesis, Inc.), by transfecting them separately at a ratio of ∼2.5E8 molecules per cell into CHO/K1 cells (see Methods). In all, there were six transfection conditions: no transfection, pTFO, nTFO, VP48, pPOC, and nPOC. We set the following cutoff threshold per gene: fragments per kilobase of exon model per million reads mapped (FPKM) value of at least 20, for at least one of the 18 samples. This resulted in 7106 of 28872 annotated genes that were included in our analysis. For each sample, we identified mRNAs that were differentially expressed (with respect to non-transfected levels and shown in the Venn diagrams in Fig 5d-f. Statistical significance was defined as false-discovery-likelihood (negative binomial test), or p-value, lower than 0.01. P-values were adjusted for multiple-testing (see Methods). In Fig. 5d, we grouped the data in accordance with classes VP48, TFOs, and POCs, and found 547, 675, and 1431 genes differentially expressed respectively. This Venn grouping shows two asymmetric features. The first is in the number of uniquely differentially expressed genes per group (25, 41, and 967 for VP48, TFOs, and POCs groups, respectively). This is consistent with the images shown in Fig. 5b-c, which show an accumulation of POCs within the cell nuclei. The second apparent asymmetry is the number of genes that are common to two of the three groups (e.g. VP48-POC group has only five genes, while the POC-TFOs and TFOs-VP48 groups have 117 and 175 genes, respectively). To further understand the origin of the asymmetry in the common genes between the groups of the differentially expressed genes, we generated another Venn diagram with four groups, this time splitting the TFOs group into the constituent nTFO and pTFO oligos (Fig. 5e). Here, the groups unique to only VP48-POCs, nTFO-POCs, and pTFO-POCs included 5, 4, and 92 genes, respectively (bold numbers in Fig. 5e), suggesting a specific regulatory interaction involving the pTFO. Finally, a Venn diagram depicting both POC and TFO subgroups reveals (Fig. 5f) that 649 of the 967 genes are differentially expressed for both pPOC and nPOC, while 266 and 52 genes of the 967 are uniquely differentially expressed for the pPOC and nPOC respectively. This indicates that while there is a large overlap between pPOC and nPOC genes, the pPOC is still associated with a statistically significant number (Fisher’s test – p-value < 2e-15) of uniquely differentially expressed genes. Consequently, the bias in the Venn diagrams toward more genes that are differentially expressed in the presence of either the pTFO oligo or pPOC provides support for a sequence dependent interaction between the oligo and some genomic targets.

Next, we hypothesized that genes regulated by a triplex interaction between the pTFO (unconjugated pTFO and conjugated pPOC) and targets on the genome would be sensitive to both low and high nuclear concentrations of oligo, with a stronger regulatory effect for higher pTFO concentrations. This interpretation assumes that the dominant regulatory role of the VP48^16^ component in the pPOC is to increase pTFO nuclear concentration. To test this hypothesis, we examined the differential expression values recorded for the 113 genes belonging to the pTFO-POC group (Fig. 5e-underline) by computing six boxplots (Fig. 5g-left) using the differential expression values recorded for the pTFO, nPOC, and pPOC transfected cells respectively, with each data-set split into positive (i.e. up-regulated) and negative (i.e. down-regulated) differentially-expressed genes. The plot shows that while there is no clear trend in upregulated genes, a clear dependence on oligo type can be observed for down-regulated genes. In particular, the increase in the magnitude of the repression when transitioning from the pTFO through nPOC to pPOC transfected cells is consistent with a thermodynamic model (see SI) for repression. In brief, the model predicts that the magnitude of the repression effect depends on the ratio between the TFO concentration to the binding affinity to its target. The model of successive stronger down-regulation effect is further supported by examining a radar plot depicting the fold-regulation values for all the genes in the 113 group (fig. 5h). For down-regulating genes (fig. 5h-left), we observe a trend where for genes with maximal fold repression values that are >1.8 (numbers around the plot - 38 of 53 genes), the pPOC generates the strongest repression effect, then the nPOC, and the weakest effect is observed for the non-conjugated pTFO oligo. By contrast such a trend only consistently appears for the top 5 up-regulated genes (fold up-regulation > 3 - fig. 5h – right).

The binding model further implies that genes that are differentially expressed only upon pPOC transfection are most likely lower-affinity pTFO targets, that respond only to high concentration of the pTFO and as a result yield a lower fold-repression level. Comparison (Fig. 5i) of the DE boxplots for the 266 genes that comprise the unique pPOC group (see Fig. 5f – bold and double underline) to those of the 113 putative high-affinity target genes that were common to the pTFO and POCs (Fig. 5e – bold and underline), we observe that the magnitude of the down-regulation is significantly stronger for the 113 as compared with the 266, consistent with our model’s predictions. Consequently, our data supports an interpretation of an oligo-dependent and thus sequence-dependent down-regulation effect (see Extended Data Fig. 7a-b for another depiction of this data using a p-value box plot).

Finally, we wanted to check if there is any correlation between the down-regulatory effect and the presence of putative TTSs in the vicinity of the down-regulated genes. To do so, we employed the triplexator calculator (http://bioinformatics.org.au/tools/triplexator/, see Methods), which computes the lengths and positions of possible TFO-TTS pairings based on canonical triplex-pairing rules. We first tested the triplexator by providing it with the N-TFO library sequences and TTS-1_37 (see Fig. 3), and plotted the triplexator scores vs our experimentally-obtained reactivity scores from the *in vitro* Triplex-Seq experiment for the same combination of TTS and TFO library (see Methods for details). The results (Extended Data Fig. 7c) show a strong correlation between the empirical reactivity and computed triplexator scores (Pearson correlation of 0.85), verifying the utility of this calculator. We next computed number-of-TTS-per-gene frequency of occurrence in the genome in a neighborhood that encompassed +/-5000 bp from the transcription start site (TSS) for each CHO gene, and plot the result in Fig. 5j (red circles and dashed lines). The results reveal an exponential distribution at the genomic level, which provides a convenient frame of reference for other populations of genes. We next computed the number-of-TTS-per-gene frequency of occurrence for the 547 VP48 genes (see Fig. 5d), and the 1084 genes in the POCs but not in the VP48 group (POCs-only and POC-TFO groups, bold numbers in Fig. 5d). We excluded the TFO-only genes (i.e. those that are not differentially expressed by the POCs), as those are not expected to be triplex-regulated according to our model. We also split each set of genes into positive and negative regulated genes. We plot the results for the down-regulated and up-regulated genes in Fig 5j-left and -right, respectively. The plots show that the number-of-TTS-per-gene frequency distribution of VP48 differentially expressed genes (green diamonds) does not deviate from the genomic distribution (red circles), for both the down- and up-regulated genes. However, for the 544 genes (of 1084) that were found to be down-regulated in POCs-only/POC-TFO group (blue squares), a clear enrichment from the genomic distribution appears for higher TTS numbers. This can be additionally observed in the inset of Fig. 5j-left, where an enriched set of datapoints appears above the eight TTSs per gene level. Conversely, for the up-regulated genes, no clear deviation is observed, consistent with the results shown both in the boxplots and radar plots (Fig. 5g-i). Consequently, our experimental and bioinformatic analysis provide evidence for a TFO specific interaction, which leads to down-regulation of genes that are enriched for TTSs in their regulatory neighborhoods.

## Discussion

In this work, we employed mixed-based oligo libraries to explore TFO sequence space and determine relative binding affinity (termed triplex reactivity) to a set of increasingly G-rich TTS sites using an approach we termed Triplex-seq. Our approach led to the characterization and verification of TFO-TTS binding rules in the form of increasingly higher information content TFM logos. Our results indicate that for triplex formation the minimal length of a TFO motif is between 7-10 nt (Figure 3), and G-rich TFO and TTS sequences (Figures 3 and 4) lead to a stronger, pH-independent, more specific triplex interaction while weak interactions between TFOs and TTS are induced using less G-rich sequences.

We then selected two variants from our various TFO libraries exhibiting both a high and low triplex reactivity potential for testing in an *in vivo* setting for a triplex-based regulatory effect. We characterized the regulatory effect of the selected TFOs using RNA-seq in both their native ssDNA form and as a synthetic biomolecule (termed POCs) where the TFOs were conjugated to a VP48 peptide in the 5’end. Our RNA-seq experiments provide two major findings. First, we observe a down-regulatory effect of x2-x4 characterized by a higher fold-repression effect for the pPOC as compared with the nPOC consistent with a higher binding affinity for the former as compared with latter in accordance with our *in vitro* findings. In addition, the dose-response-like behavior observed for 48 of 53 genes that were both down-regulated for the pTFO and pPOC (Fig. 5h-left), and the appearance of 266 unique pPOC genes exhibiting lower repression are all consistent with a sequence specific interaction between the pTFO and some genomic targets. The second major finding is a modest enrichment in putative matching TTS loci only in the genomic neighborhoods of POC down-regulated genes.

What could be the mechanism underlying the down-regulatory effect? It would seem that the most likely explanation involves binding of the POC constructs via a triplex interaction to open chromatin in the vicinity of active genes, which likely interferes with the binding of transcriptional machinery (e.g. by inhibiting transcription factor binding^15,18,48^) leading to the observed down-regulatory effect. While we do not provide definitive proof for triplex interaction *in vivo*, there are not many alternative sequence-specific interactions that can be offered to explain our data (e.g. ssDNA binding to RNA thus triggering degradation^49^). This is further compounded by the fact our TFOs are ssDNA oligos, which in turn exclude popular RNA-based alternative explanations to triplex formation such as an interaction with some putative RNA-DNA binding protein. Consequently, triplex formation *in vivo* is at the very least a reasonable explanation for the observed down-regulatory effect.

Finally, irrespective of the underlying mechanism we provide here a new molecular tool for genomic manipulations. While native ssDNA oligos are not known to exist within cells for an extended duration of time, thus severely affecting their utility, the conjugated form presented here was found to exist stably within cell nuclei for at least 24 hrs using confocal microscopy. The retention affect also manifested itself in 1431 differentially expressed genes for the pPOC experiment as compared with 557 genes for the pTFO one. This, therefore, suggests that conjugation of an ssDNA oligo to the VP48 peptide facilitates not only entry into the nucleus and contributes to its retention, but also leads to enhanced regulatory activity. In addition, the peptide-oligo conjugate does not require an *a priori* transfection or infection of a large protein (e.g. Cas9) to unleash its regulatory function, and can thus play an important technological role in non-transfectable or difficult-to-transfect cell lines. It is important to note that the down-regulation effect observed is contrary to our initial design of the VP48 peptide that was intended to activate expression. Thus, even though the peptide part of the POC studied here does not seem to exhibit its intended functionality, the resultant specific regulatory effect was still sufficiently potent to generate widespread disruption in gene expression (for ∼1000 of the 7000 genes that were significantly detected). Given the fact that the POCs cannot be genetically encoded, future research and development of this and other novel synthetic biomolecules may provide an important new avenue in genome regulation and editing research, and related therapeutic approaches.

## Methods

### Design of triplex-forming oligos (TFOs) and triplex-target site (TTS)

The developed Triplex-Seq assay requires oligonucleotides (oligos) which are referred to as triplex-forming oligos (TFOs) and triplex target sites (TTS) and were ordered from Integrated DNA technologies (IDT) as desalted oligos. The double-stranded TTS were generated by annealing complementary oligos, whereas the single-stranded TFO libraries were synthesized using the standard mixed bases tool from IDT. Details about the oligos can be found in the Supplementary Methods.

### Triplex formation *in vitro*

To trigger triplex formation *in vitro*, 1000 pmole of TFOs and 50 pmole of a respective TTS were mixed at a molar ratio of 20:1 (TFO:TTS) and incubated in appropriate buffer conditions (pH 7/anti-parallel: 10 mM Tris-HCl pH7.2, 10 mM MgCl_2_; pH 5/parallel: 10 mM sodium acetate pH 5.0, 10 mM MgCl_2,_ triplex-disfavoring: 10 mM Tris-HCl pH7.2, 10 mM MgCl_2_, 140 mM KCl) at 37 °C for 2 hours in a final volume of 25 µL. Samples were either subjected to the DNA ScreenTape assay (2200 Tapestation, Agilent) using 1 µL of each sample or mixed with 1x DNA loading dye (NEB) and loaded on a 10 % native polyacrylamide gel (PAGE) for separation of TFO, duplex and triplex fragments (see more details in description of electrophoretic mobility shift assay below).

### Electrophoretic mobility shift assay (EMSA)

To separate triplexes from duplex DNA and non-bound TFOs, a native polyacrylamide gel (PAGE) was used. The 10 % -15 % PAGE was prepared by polymerizing the acrylamide/bis-acrylamide 40 % solution (Sigma) using N,N,N’,N’-Tetramethylethylenediamine (Alfa Aesar) and ammonium persulfate (Sigma) in respective buffers (anti-parallel, parallel and triplex-disfavoring buffers). Following PAGE preparation, samples were mixed with 1x purple loading dye (6x, NEB) and 7.5 µL of low molecular weight DNA ladder (NEB, #N3233) was loaded onto the gel. The 1x running buffer is the same that has been used for PAGE preparation. For sufficient band separation between triplexes and duplex DNA, electrophoresis was operated for 2 hours at a field strength of 7.5 V/cm^2^. Subsequently, the gel was removed from the electrophoresis chamber and transferred to 1x running buffer containing 0.1 mg/mL of ethidium bromide (1 mg/mL, Hylabs) to stain DNA for 20 minutes at RT while carefully shaking. Images of gels were acquired using a UV gel documentation system.

### DNA fragment isolation from PAGE

Following triplex/duplex/TFO separation, DNA was isolated from PAGE using the Crush and Soak Method described by J. Sambrook and D. Rusell. In brief, while UV illuminating the PAGE (305 nm), putative triplex bands in triplex, duplex, or TFO-only lanes were excised at the same height corresponding to the triplex DNA from the positive control in the triplex sample using a clean scalpel. Extracted gel slices were transferred into 1.5 mL microcentrifuge tubes. The weight of the slice was determined and 2 volumes of 1x Crush and Soak buffer (CSB, 200 mM NaCl, 10 mM Tris-HCl pH7.5, 1 mM EDTA pH8.0) were added. The gel was crushed into smaller fragments using a sterile pipette tip or inoculation loop and incubated overnight at 37 °C while slowly shaking. Following the overnight incubation, samples were centrifuged at maximum speed (16000 *g*) for 2 minutes at 4 °C. The supernatant was transferred to a fresh microcentrifuge tube and an additional 2 volumes of CSB were added to the gel pellet, centrifuged (16000 *g*, 2 minutes, 4 °C) and supernatants were pooled. Subsequently, DNA was ethanol-precipitated by addition of 3 volumes of ice-cold ethanol, 1/10 of volumes sodium acetate (pH5.0) and 1 µg GlycoBlue Coprecipitant (glycoblue, 15 mg/mL, Thermo Fischer Scientific). Samples were incubated for at least 1 hour at -80 °C followed by centrifugation (16000 g, 30 minutes, 4 °C). Supernatant was carefully decanted, DNA was air-dried for 5 minutes at room temperature and dissolved in 15 µL of ultra-pure water (Ultra Pure Water, Biological Industries).

### Heat-separation of duplex and TFO DNA (triplex disruption)

To ensure that TFOs are not bound to duplex DNA, which is required for the ssDNA adapter ligation in the next step, the DNA from the previous steps was mixed with 1x triplex-disfavoring buffer (TDB: 10 mM Tris-HCl pH7.5, 140 mM KCl) and incubated at 95 °C for 5 minutes to separate duplex DNA from the TFOs. Subsequently, DNA was reannealed gradually by decreasing temperature by 1 °C every 30 seconds until room temperature was reached. Following reannealing of duplex DNA and simultaneous prevention of triplex formation, DNA was ethanol-precipitated as has been described above (3 volumes of ice-cold ethanol, 1/10 of volumes sodium acetate (pH5.0) and 1 µg GlycoBlue) and resuspended in 23 µL ultra-pure water.

### Single-stranded adapter ligation

After TFOs were separated from duplex DNA, ssDNA adapter ligation was performed using the CircLigase ssDNA ligase kit (#CL4115K, Epicentre). Briefly, samples were mixed in 1x CircLigase buffer, 2.5 mM MgCl_2_, 50 µM adenosine-triphosphate (ATP), 100 U CircLigase and 50 pmole ssDNA adapter which contains a 5’ phosphorylated terminus (to act as donor) and a 3’ carbon spacer (for more details see Table 5). The reaction mix was incubated for 2 hours at 60 °C with subsequent deactivation of the enzyme for 10 minutes at 80 °C. The obtained TFO fragments were purified using Agencourt AMPure beads (Coulter Beckman), according to manufacturer’s instructions. In brief, 1.8x volumes well-resuspended AMPure XP bead slurry was added to the PCR reaction mix, incubated for 5 minutes at RT and transferred to the DynaMag-96 Side Magnet (#12331D, Thermo Fisher Scientific). Following incubation of the sample on the magnet for 2 minutes (or until sample is clear), supernatant was removed and beads were washed twice with 200 µL freshly-prepared 70 % ethanol without removing samples from the magnet. Subsequently, samples were removed from plate, air-dried for 5 minutes to ensure no residual ethanol was left, resuspended in 25 µL ultra-pure water and incubated for 5 minutes at RT before transferring to magnet. Following a 2 minutes incubation, supernatant was carefully transferred to a fresh tube.

### Preparation of sequencing library

The final step of the protocol is the PCR amplification of the enriched TFOs and simultaneous addition of Illumina adapter sequences including indexes to multiplex samples. For a detailed list of Illumina oligonucleotides, index sequences and other PCR primers see Table 5. For the PCR mix, 0.01 µM of primer TriSeqNGS001 which binds the capture sequence of the TFO, 0.5 µM Illumina primer index #1-#34 that bind the ligated ssDNA adapter sequence and adds Illumina indexes, 200 µM dNTPs (each dNTP 100 mM Solution, Thermo Fisher Scientific), 1 U Q5 Hot Start High-Fidelity Polymerase (Q5, NEB), 3 % dimethylsulfoxide (DMSO) were mixed in 1x Q5 Reaction buffer and the following PCR program was executed: Initial denaturation for 2 minutes at 98 °C, followed by 15 cycles of 30 seconds at 98 °C, 30 seconds at 65 °C and 10 seconds at 72 °C, which preceded the final elongation step for 2 minutes at 72 °C. PCR samples were purified using AMPure XP beads as has been described above. In a second PCR, 0.5 µM Illumina primer index #1-#34, µM primer PE_forward, which adds the sequence that is complementary to the Illumina flow cell, were added to the reaction mix as has been described above. The same PCR program was used and after PCR completion, samples were cooled down to 4 °C, 5 U of Exonuclease I (ExoI, NEB) were added to the PCR mix and incubated at 37 °C for 30 minutes. Samples were subsequently purified using AMPure XP beads as described above. DNA Screen TapeAssay and Illumina sequencing 1 µL of prepared double-stranded DNA (dsDNA) libraries were analyzed by the DNA ScreeTape assay and size of DNA fragments was determined. To multiplex and prepare sequencing libraries, the molarity of the PCR-amplified dsDNA libraries was calculated by determining the average length based on the Tapestation results and the concentration of the dsDNA fragment which was measured by Qubit 4 Fluorometer (Thermo Fisher Scientific) and samples were to obtain a 10 nM pooled and multiplexed library.

### Illumina sequencing

The multiplexed libraries were sequenced either on an Illumina HiSeq 2500 (High Output Run Mode V4 or Rapid Run Mode) as a single run or as a spike-in which included the addition of 5-10 % of the pooled library to another prepared library (operated at the Technion Genome Center, Haifa) depending on the number of different variants of the pooled library as a single-read 50 bp run. Due to the low diversity of sequences in the libraries, we added 20 % PhiX Control v3 Library (Illumina, FC-110-3001). The overall read yield ranged between 150 - 300 million reads per HiSeq run.

### Cell culture

The cell line CHO/K1 (kindly provided by Arie Admon’s lab, Technion) was incubated and maintained in 100×20 mm cell culture dishes (Nunclon cell culture treated, Thermo Scientific) under standard cell culture conditions at 37 °C in humidified atmosphere containing 5% CO_2_ and were passaged at 80-85 % confluence. Cells were washed once with 1x DPBS (Biological Industries), and subsequently treated with 1 mL trypsin/EDTA (Biological Industries) followed by incubation at 37 °C for 1-2 minutes. DMEMcomplete (Dulbecco Eagle’s Minimum Essential Medium, Biological Industries), complemented with 10% FBS (Biological Industries, Lot.no: 1418110) and final concentrations of 100 U penicillin plus 100 μg streptomycin (Biological Industries), was added and transferred into fresh DMEMcomplete in subcultivation ratios of 1:10.

### Peptide-oligo conjugates (POCs) synthesis

Both custom POCs (as well as unconjugated VP48 peptide) were produced by Bio Synthesis, Inc. The VP48 was conjugated separately to the 5’ ends of pTFO and nTFO oligos through a succimidyl 4-(N-maleimidomethyl) cyclohexane-1-carboxylate (SMCC) hetero-bifunctional linker. The nTFO POC had an AlexaFluor488 fluorophore added on the 3’ end of the nTFO. Both POCs were HPLC-purified and subjected to mass-spectrometry by the manufacturer, and had POC purities of over 90%.

### Transient transfection

CHO/K1 cells were seeded at a density of 1 million cells per plate or 20k cells per plate on 10cm and 35mm plates, respectfully, and incubated at 37°C. 24h post seeding, cells were transfected using polyethylenimine (PEI, from Polysciences, Inc. cat. no. 23966-1) in OptiMEM (Gibco, cat. no. 31985070). PEI stock concentration was 1 mg/ml, in ultrapure water. Molar ratios of ∼4.5E-16 and ∼1.1E-12 DNA/cell and PEI/cell, respectively, were maintained across transfections. After 24 hours, at least 1E6 cells from the 6 transfection conditions were pelleted and stored at –80 °C. There were three replicates for each experimental condition, resulting in 18 samples. RNA extraction, library preparation, and sequencing were performed on all samples.

### Microscopy

24h post transfection, CHO/K1 cells were incubated in PBS with 0.1% Hoechst stock solution (Hoechst stock concentration: 5 mg bisBenzimide H 33342 trihydrochloride, Sigma Aldrich cat. no. B2261, in 1 ml ultrapure water) and incubated at 37°C for 10 min. Then, cells were washed twice with PBS, and imaged by a Zeiss LSM 700 inverted confocal microscope in PBS with 5% fetal bovine serum (FBS, Biological Industries, cat. no. 04-007-1A).

### RNAseq

Three replicate samples were prepared for each experimental condition. Samples were prepared for RNAseq following an online protocol (https://www.activemotif.com/). Each sample contained approximately 1 million cells. Cells were rinsed once with PBS to remove dead/detached cells, subjected to 0.05% trypsin in 10 ml PBS, and transferred to a 15 ml tube. Cells were centrifuged at 800 g for 5 min at 4°C, after which the supernatant was removed. Cells were then resuspended by vortex in 10 ml ice-cold PBS and subjected to 800 g for 5 min at 4°C, after which the supernatant was removed. Cells were then resuspended in 1 ml ice-cold PBS by vortex and transferred to 1.5 ml tubes. Finally, cells were centrifuged for 5 min at 800 g and 4°C, supernatant was removed, and pellets were flash-frozen in liquid nitrogen. Samples were stored at –80 °C (upright freezer) and transferred on dry ice to the Technion Biomedical Core Facility (BCF). There, RNA was extracted from cells using the RNeasy mini kit (Qiagen, cat no. 74106) and the Qiacube automated system (Qiagen). Quality control for total RNA was performed using TapeStation (Agilent). The RINe value of all samples was > 9.7. RNAseq libraries were produced from all samples according to the manufacture protocol (NEBNext UltraII Directional RNA Library Prep Kit for Illumina, cat no. E7760), using 800 ng total RNA per sample. mRNA pull-up (polyA selection) was performed using Magnetic Isolation Module (NEB, cat no. E7490) and sheared before ligation of library primers. All 18 libraries were mixed into a single tube, with equal molarity. The RNAseq data was generated on an Illumina NextSeq500 75 SE high-output flow-cell (Illumina, cat no. FC-404-2005). The run yielded approximately 550 million reads, with ∼30M reads per sample. The cluster density was ∼230 with ∼90% cluster passed filter.

### Bioinformatic analysis

#### Post-sequencing processing

For every TFO-TTS Triplex-Seq experiment, we analyzed the Illumina library pair of the TFO-only and triplex bands as follows: first, Illumina sequencing read quality was validated, adapter sequences were trimmed using cutadapt^50^ and aligned to the PhiX genome bowtie2^51^ in local alignment mode (bowtie2 --local). Second, TFO sequences were extracted by (i) identifying the 19 nt long capture sequence ‘CTTCAGCTTGGCGGTCTGG’, (ii) selecting sequences with exactly 39 nt and (iii) searching for identical matches to all possible combinations of the TFO sequence. Next, the number of reads were normalized by dividing each read count by the total number of reads of the triplex lane and multiplied by 10^6^ (reads per million, RPM). The frequency of these RPM values were plotted as a function of RPM (frequency plot). Finally, for every variant in the TFO library, the triplex reactivity was calculated. Triplex reactivity is defined as the ratio of the RPM of the triplex band divided by the RPM of the TFO only band and subtraction of 1. Thus, every value above 0 is defined as “triplex reactive”:

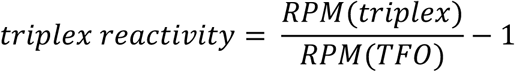

#### Nucleotide frequency computation

To obtain insight into the distribution of nucleotides within an enriched TFO at a given triplex reactivity score, the frequency of each nucleotide (G, A, T, C) in the 20 nt long triplex-forming sequence was computed. The average of the frequencies of TFO sequences with the same triplex reactivity value was calculated and plotted as a function of triplex reactivity values for each TFO library.

#### DRIMust analysis and k-mer density and positioning computation

DRIMust (Discovering Ranked Imbalanced Motifs using suffix trees) is a tool to compute enriched k-mers and motifs based on a ranked list of sequences. Here, the triplex reactivity lists of each TFO library were sorted (from highest to lowest triplex reactivity value), the first 40,000 (if applicable) lines were converted to a fasta file and uploaded on the DRIMust website (http://drimust.technion.ac.il/index.html). The parameters for the DRIMust motif and k-mer computation were set as follows: Motif length range: 5-20 nt, statistical significance threshold: 10^−6^, maximum number of motifs to display: all motifs. Following the computation, we obtained a list of k-mers (short motifs with a length that range between 5-20 nt) and a sequence logo (DRIMust motif). These motifs and k-mers are shown throughout the manuscript. To further evaluate the occurrence of k-mers within the enriched TFO sequences, the running average of the density of each k-mer that was identified in the enriched TFO sequences for a 100 variant window was calculated. Finally, this density was plotted as a function of number of variants. To characterize the position of k-mers within the enriched TFO sequences, the start position of each detected k-mer was determined and the number of k-mers at a given start position was plotted as a function of start position.

#### Differential expression analysis pipeline

The NGS reads for each sample were processed using a custom bash shell script, as follows. All sample reads were combined into one file, after which functions `fastqc` () and `trinnomatic` () were applied. This resulted in 25-31 million reads per sample.

Our reference genome was the file cgr_ref_CriGri_1.0_unplaced.fa For gene annotations, we used a modified version of the annotation file GCF_000223135.1_CriGri_1.0_genomic.gff, from which we removed all tRNA, rRNA and miRNA. (Both files were downloaded from https://blast.chogenome.org/archive/Genomes/.) We prepared the reference for alignment using `rsem` (http://deweylab.github.io/RSEM/rsem-prepare-reference.html). We then mapped all RNAseq sample reads to the annotated reference genome using `rsem` (http://deweylab.biostat.wisc.edu/rsem/rsem-calculate-expression.html). This yielded the raw number of counts and the Fragments Per Kilobase Million (FPKM) values for each sample.

We checked the degree of sample similarity within and between experimental triplicates using cross-correlation (see Extended Data Fig. 8). We then calculated differential expression (DE) for each experimental condition relative to the non-transfected experimental control using custom Matlab code, as follows. We first identified genes with maximal FPKM value across all samples lower than 20 and excluded these genes from DE analysis. We then calculated the likelihood that expression levels of each transfection condition differ from expression levels of the non-transfected control using Matlab function `nbintest` with options ‘VarianceLink’ and ‘Identity’. Adjusted fals^52^e discovery rates under the null hypothesis (namely, no change in expression) were calculated using Matlab function `mafdr` with the ‘BHFDR’ (Benjamini-Hochberg) option on the output of `nbintest`. We refer to these adjusted false discovery rates as p-values throughout the main text. Fold-change was calculated by dividing the mean FPKM values for each transfection condition by the mean FPKM values of the non-transfected control. A gene is considered to be differentially-expressed (DE) if it satisfies p-value < 0.01. The Venn diagrams displaying the DE genes of all subsets of the 5 transfection conditions relative to the non-transfected control was generated using http://www.interactivenn.net/^52^. See Supp. Table 1 for all differentially expressed genes and relevant sub-groups.

#### TFO scoring and Triplexator analysis

The Triplexator VM image was downloaded from http://bioinformatics.org.au/tools/triplexator/inspector/install.html and run on a Virtual Box on a Mac desktop station running macOS High Sierra (v10.13.6). This prediction software Triplexator^53^ analyzes potential TFO-TTS pairs by matching the ssDNA to the dsDNA applying user-specific parameters. This powerful, computational framework also predicts putative TFOs within an ssDNA sequences, or potential TTS in dsDNA. The main parameters that were used to generate TFOs matching the TTS with increasing percentages of guanines are listed in below:

- maximum error-rate: 10%; maximum total error : 3; maximum number of tolerated consecutive pyrimidine interruptions in a target: 1
- minimum guanine content with respect to the target : 10% ; maximum guanine content with respect to the target : 100%
- minimum length : 10 nt- maximum length : 30 nt
- minimum guanine-percentage in anti-parallel mixed motif TFOs : 0% ; maximum guanine-percentage in parallel mixed motif TFOs : 100%

#### TTS identification within genomic regions using triplexator

We defined candidate TTSs as genomic sequences of length 10 bp or longer with a triplexator score of 10-13 with a guanine percentage of at least 80%. Furthermore, we confined our search to nearby the promoter region at distances of +/- 10000 bp from the transcription start site of the RNA sequences detected. Number-of-TTS-per-gene frequency Analysis was confined to +/- 5000 bp. See Supp. Table 2 for all TTS identified within +/- 10000 bp for all TSSs.

#### Thermodynamic model for TFO-based repression

Repression can be modelled as a 2-state thermodynamic model^54^. Here we assume that there are two states: DNA bound by protein, and DNA unbound. In the case of a repressor, the unbound state is the transcriptionally active state, which can in steady state leads to the following relationship between the target gene and repressor concentrations:

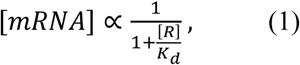

where *[mRNA], [R]*, and *K*_*d*_ correspond to the concentration of target gene mRNA, the concentration of the repressor, and its binding affinity to the target site respectively. Since differential expression was defined as the ratio between the mRNA concentration in the transfected experiments to the non-transfected control, we can now express this quantity using the following equation:

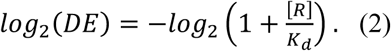

In the case of our system, the repressors are nTFO, pTFO, nPOC, and pPOC. Based on the microscopy experiments we can make the following statement:

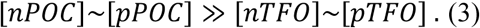

Moreover, since the pTFO is assumed to be more specific, we can also assume that:

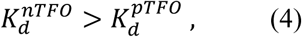

indicating that the pTFO is has a stronger binding affinity to the dark side. This implies the following:

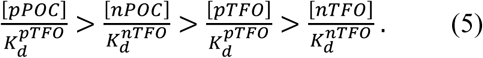

Putting everything together, we can now make the following prediction for the differential expression for any particular gene containing a TTS cluster matching the pTFO:

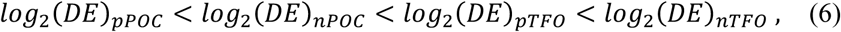

which is precisely what was observed in the experiments.

## Data availability

The datasets generated during and/or analyzed during the current study are available from the corresponding author upon request.

## Acknowledgements

Acknowledgments are also given to Inbal Vaknin, Michal Brunwasser, Naor Granik, Noa Katz, Nanami Kikuchi, Or Willinger and Roni Cohen for discussions. Ben-Zion Levi, Arnon Henn, and Avi Shpigelman (all from Technion – Israel Institute of Technology) are acknowledged for support with materials.

## Author contributions

B.K. designed and carried out the experiments, and performed the analysis of the data. O.W. carried out the RNA-seq experiments and analysis. N.E. carried out some of the experiments. L.K. implemented the initial platform. L.A. assisted implemented the triplexator analysis. O.S. assisted with the bioinformatics analysis. O.A. assisted with some of the experiments. S.G. assisted with the RNA-seq experiments, bioinformatic and triplexator analysis. R.A. and Z.Y. supervised the study. B.K. and R.A. wrote the manuscript.

