## Extended Data for "Identifying triplex binding rules *in vitro* leads to creation of a new synthetic regulatory tool *in vivo*"

### Extended Data Figure 1

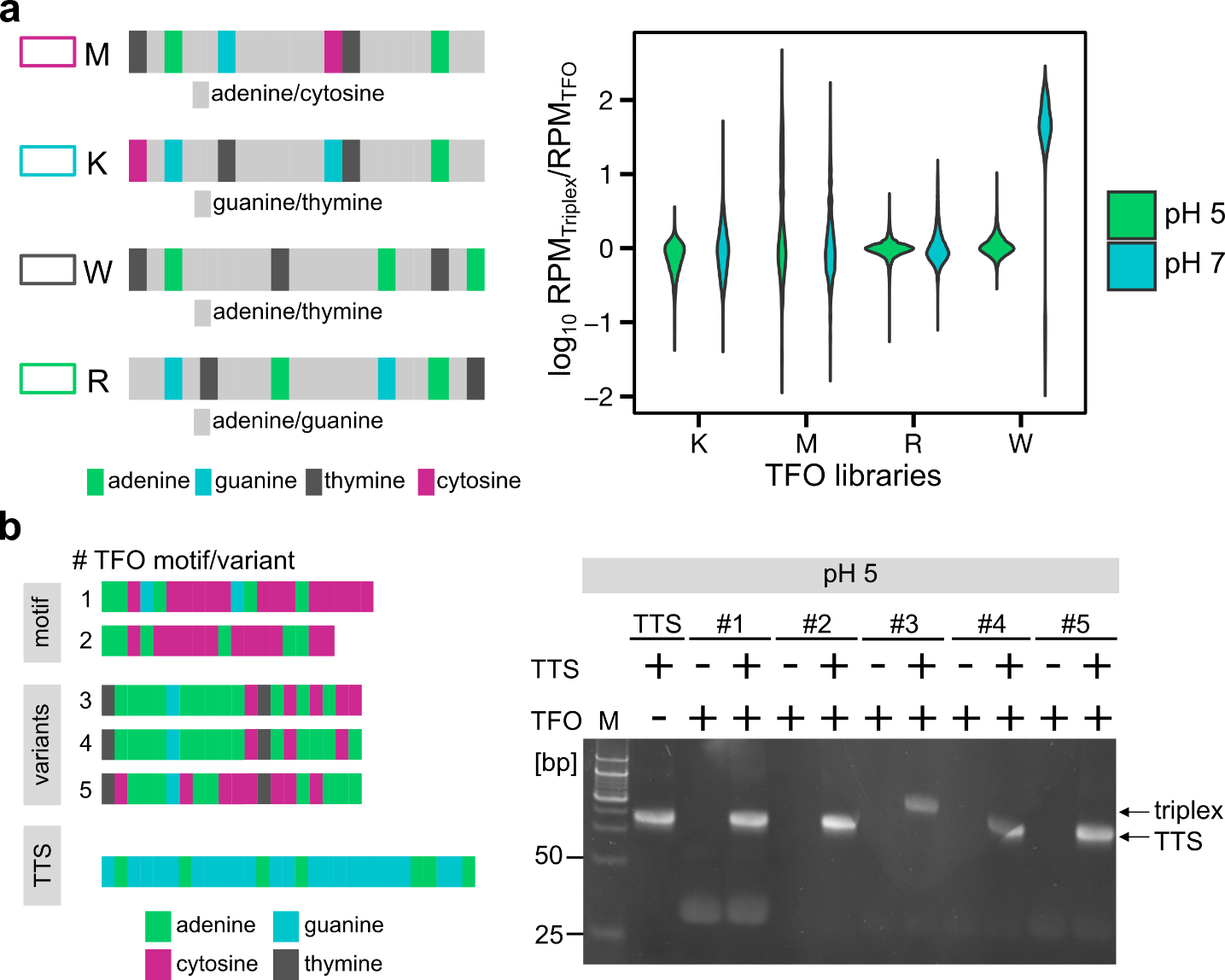

**Extended Data Figure 1| Distribution of triplex reactivity scores and variants tested for triplex potential. a, The ratio of RPM_triplex_ and RPM_TFO_ of four 16,000-variant large TFO libraries tested in pH 5 and pH 7 condition are shown in the violin plot. An acidic environment (pH 5, green) results in overall lower triplex reactivity values compared to neutral pH (pH 7, blue). b, Verification experiments for single TFOs identified by Triplex-Seq.** Two TFO motifs (#1 and #2) based on the DRIMust logo (see Figure 2c, right panel) were chosen, sequence was duplicated and ordered as a double repeat to further increase the possibility for triplex formation. Three variants with high triplex reactivities (#3-5) were selected from the sorted triplex reactivity list and tested for their potential to form triplexes in pH 5. As can be seen in the 15 % PAGE, only one TFO variant (#3, lane 8) shows a triplex band, while the other lanes lack the triplex bands.

### Extended Data Figure 2

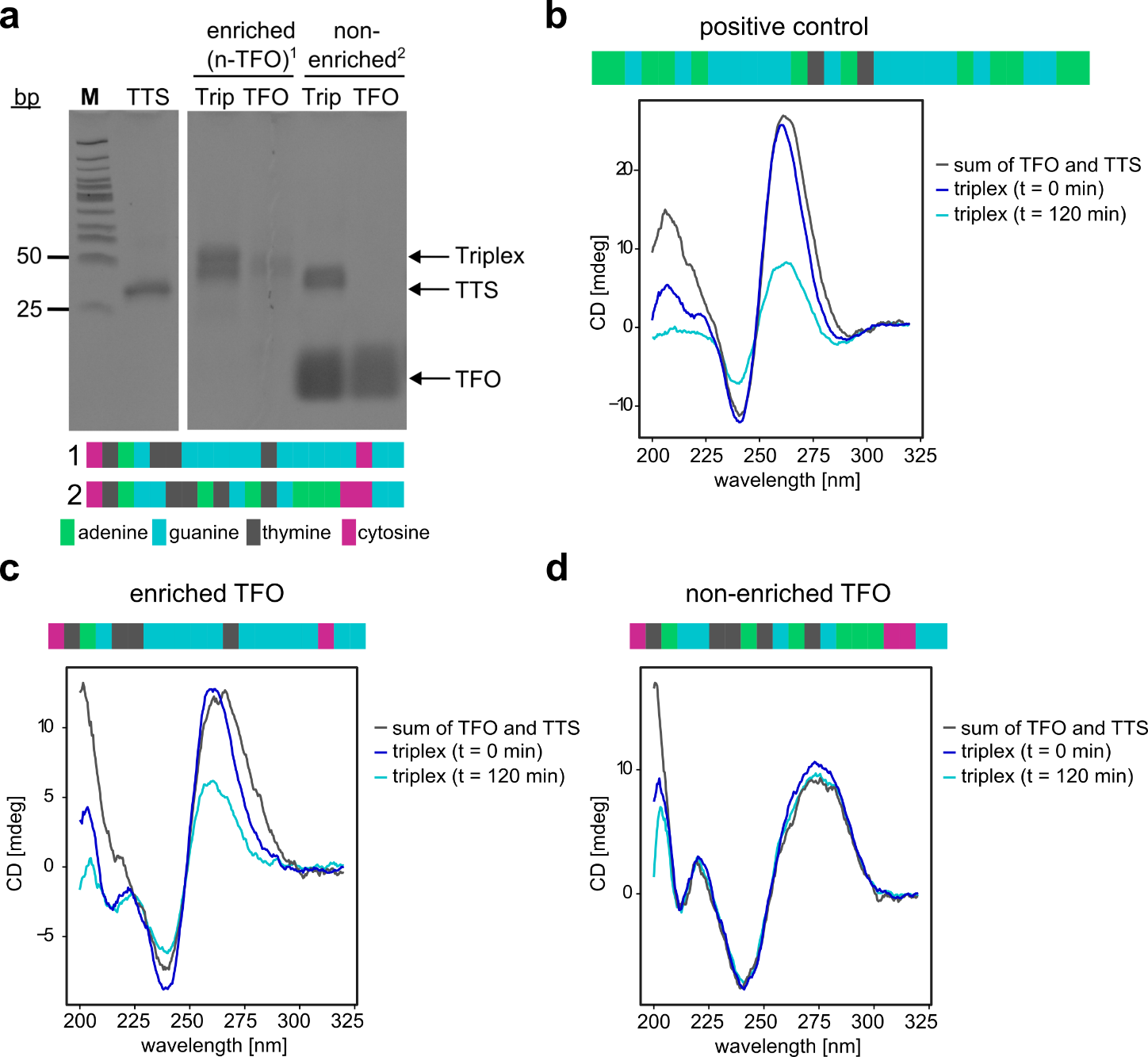

**Extended Data Figure 2| EMSA and circular dichroism spectroscopy to verify triplex forming potential of N-TFO library. a,** An electrophoretic mobility shift assay (EMSA) was performed on a highly-enriched TFO variant (highest triplex reactivity score) and compared to a non-enriched variant (negative triplex reactivity score) and a clear shift for the enriched N-TFO variant is observed, contrary to the non-enriched N-TFO variant where no shift is noted. **b-d,** Circular dichroism (CD) spectroscopy of three different samples. CD spectroscopy is a technique to discriminate between DNA structures by the differential absorption of left- and right-handed circularly polarized light and the difference between the absorption is plotted as CD in millidegrees [mdeg]. **b,** The AG30 positive control shows a difference in absorption between the sample that was measured immediately after mixing (t=0 min) and after incubation in which triplexes had time to form (triplex sample). A reduced, positive CD peak at 260 nm and a flat, slightly negative absorption is observed for the triplex sample at 205 nm. No difference between the CD spectra directly after mixing and the CD spectra for the sum of the individual spectra for the TFO and the TTS is observed. **c,** Similar CD spectra are shown where the enriched N-TFO variant was tested and the most notable change is based on the slightly skewed CD curve with a peak at 260 nm which is only observed for the triplex sample (t=120 min). **d,** No differential absorption of all three samples is observed for the non-enriched TFO variant.

### Extended Data Figure 3

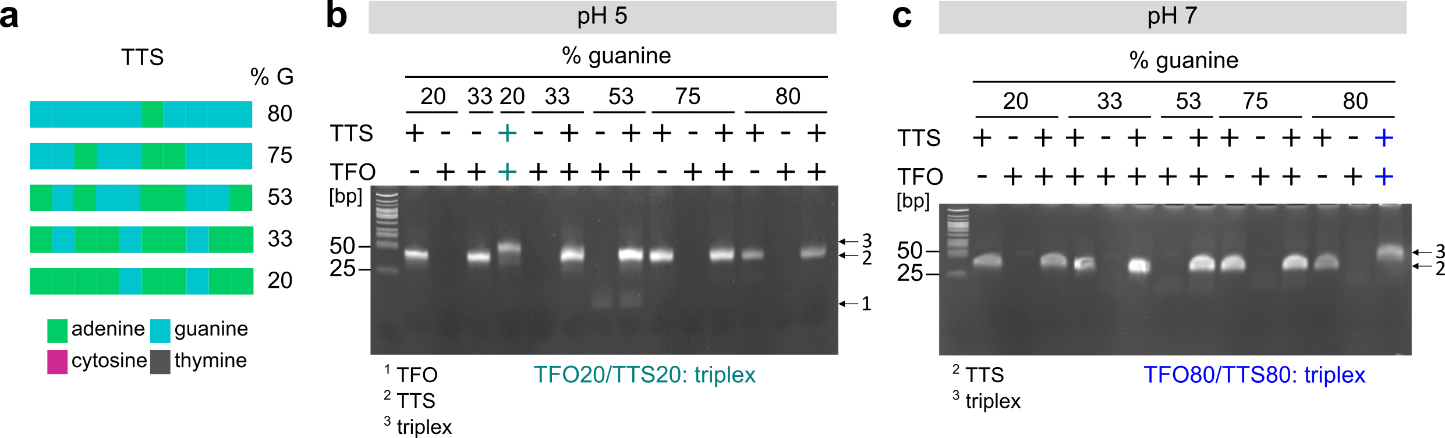

**Extended Data Figure 3|** **Mobility shift assay to identify *de novo* TFO/TTS pairs.** An electrophoretic mobility shift assay (EMSA) was performed on the five *de novo* designed TTSs with increasing guanine content. **a,** TFOs (see Supplementary Table 1) for every TTS were generated and triplex formation potential was predicted using the Triplexator software. Each TFO/TTS pair (e.g. the TFO20 was tested with the TTS with 20 % guanine) was tested in pH 5 and pH 7 conditions. **b,** The PAGE in the pH 5 condition identifies one working TFO/TTS pair that exhibits a shift from duplex to triplex (lane 4, highlighted in green) for the TTS with 20 % guanine. **c,** In the PAGE that was tested in the pH 7 condition, also one TFO/TTS pair was identified that forms triplexes (lane 14), which corresponds to the TTS with 80 % guanine. These TFO/TTS pairs were used as positive controls in the subsequent experiment in which the five different TTS variants were used.

### Extended Data Figure 4

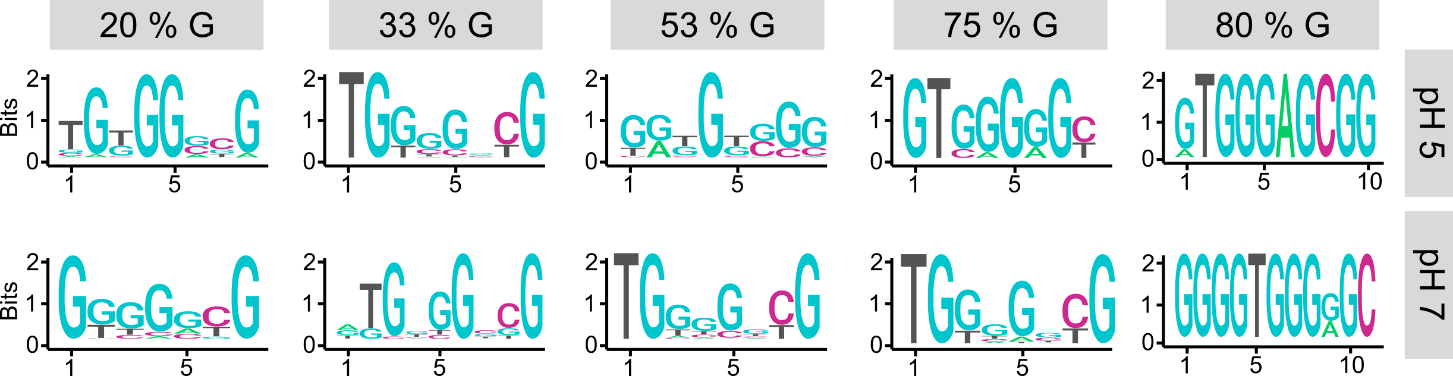

**Extended Data Figure 4|** **G-rich TFO consensus motifs identified with de novo designed TTS variants**. A DRIMust analysis was used to characterize consensus motifs for the N-TFO library that was tested with each TTS (identified by the percentage of G in each column) in pH 5 (upper row) and pH 7 (lower row). It can be noted that guanine is dominating in all consensus motifs, but little information can be retrieved for TFOs with low to medium guanine content in the TTS (20-50% G) suggesting a weaker and less-specific triplex interaction of the TFOs with the TTS variants. This is in contrast to the TTS with high guanine content (in particular for 80 % G) only one or two motifs dominate in the TFO sequences implying a strong and specific interaction with TTS variants with high guanine content (p-values between 8x10^-60^ and 5x10^-324^).

### Extended Data Figure 5

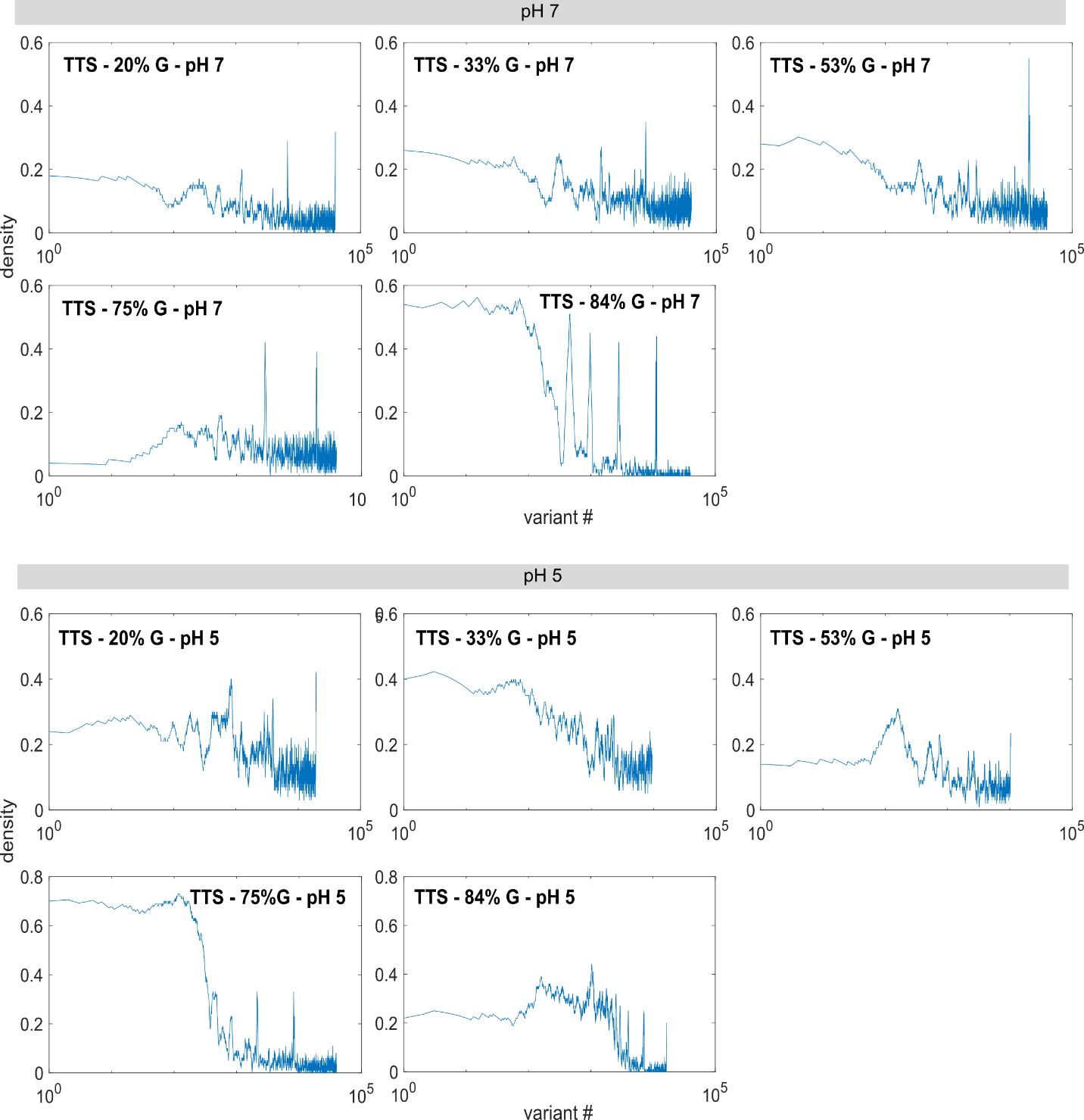

**Extended Data Figure 5| Density plots of k-mers of N-TFO library with TTS variants that differ in guanine content.** The density of k-mers that were identified within the enriched TFO sequences were plotted as a running average of a 100 variant window as a function of variant number. As shown in the plots, a sharp step function for G-rich TTS (75-80 % guanine) in pH 5 and pH 7 is observed with intermittent peaks which is particularly visible for the TTS variant containing 80 % guanine in pH 7 (upper left). Variants with lower guanine content (20-50 %) in pH 5 and pH 5 and pH 7 display a gradual decline of k-mer densities with an increasing number of variants.

**Extended Data Figure 6**

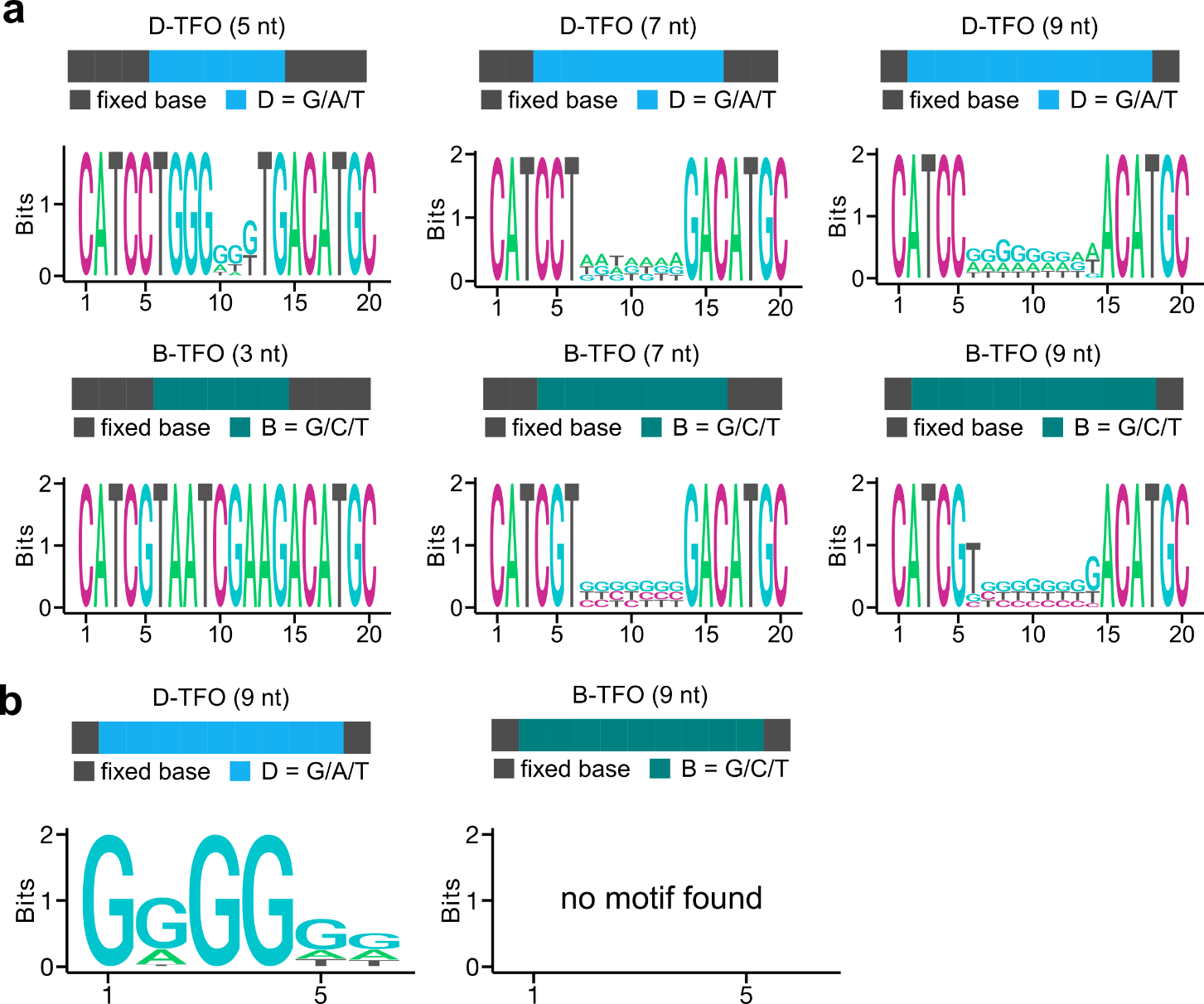

**Extended Data Figure 6| Mixed-base stretches *in vitro* identify minimal TFO length.** To characterize a potential minimal length of a TFO that is required to form stable and specific triplexes, TFO libraries with mixed-base stretches (B- and D-TFO) of varying length (3-9 nt) flanked by fixed bases were designed and tested in pH 5 (B-TFO) and pH 7 (D-TFO). **a,** The sequence logos of the D-TFO with 5 nt (left), 7 nt (middle) and 9 nt (right) are shown. The 3 nt D-TFO library is not shown as no sequence reads were obtained. Only for the 5 nt and the 9 nt long D-TFO stretch, a trend towards G-rich stretches can be observed. The sequence logos for 3 nt (left), 7 nt (middle) and 9 nt (right) for the B-TFO are displayed. While only one TFO variant dominated in the 3 nt TFO library, a trend to G-rich sequences can be seen for the 7 nt and 9 nt long B-TFO stretches. **b,** To identify a consensus motif, DRIMust was applied and only in the D-TFO (9 nt) library a G-rich stretch of 5 nt was found, while for the other libraries no DRIMust motif was identified.

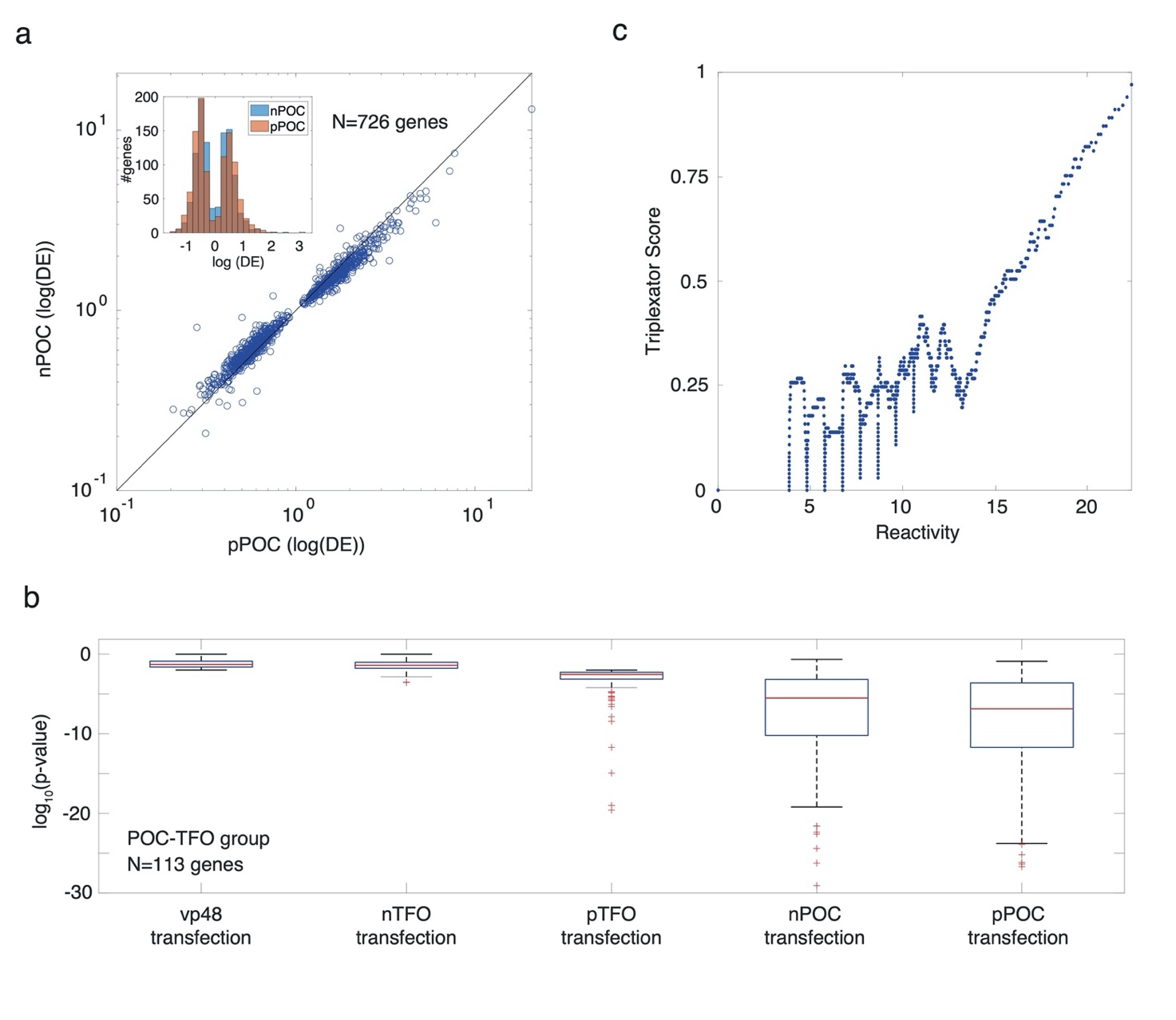

**Extended data figure 7.** (a) Plot depicting the increased regulatory effect of pPOC as compared with nPOC for the 726 genes comprising the pPOC-nPOC and pPOC-nPOC-pTFO groups in Fig. 5F. (b) p-value distributions for the 113 POC-TFO genes obtained for all 5 transfection experiments. Genes were selected to be differentially expressed if the p-value<0.01. Here the p-value boxplots reflect the differential expression magnitude shown in Fig. 5h, and provide additional support to the thermodynamic model for regulation by showing a similar trend for the p-values as for the differential expression values. Plot showing the computed triplexator TFO score for the n-TFO library, as a function of the empirical reactivity score.

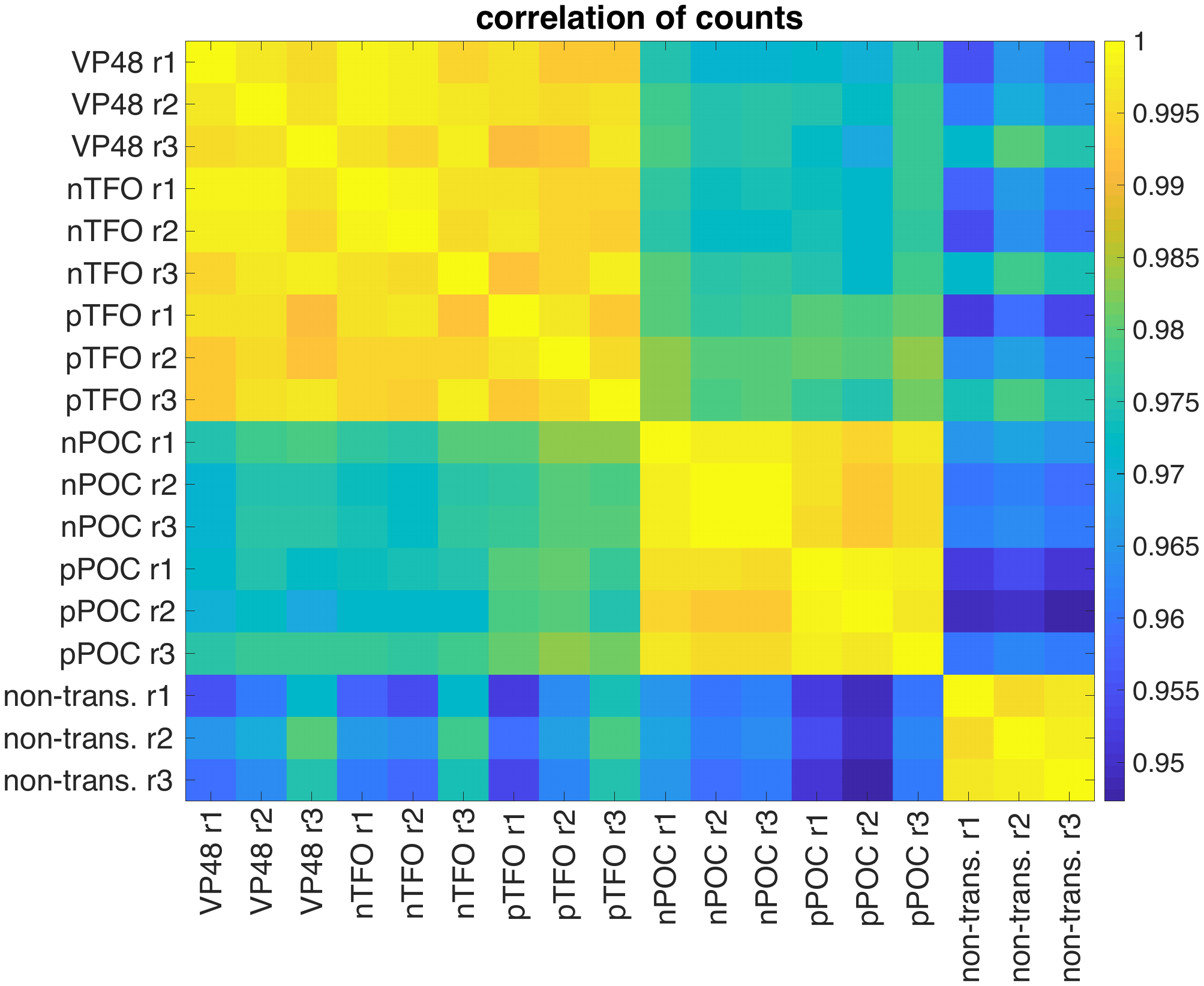

**Extended data figure 8.** Pairwise linear correlation coefficients of gene counts for each 2 RNAseq samples. ri, i=1,2,3 indicate replicates of the same experimental condition.
