## Supplementary Methods for "Identifying triplex binding rules *in vitro* leads to creation of a new synthetic regulatory tool *in vivo*"

Supplementary Information

### Supplementary Methods

#### Design of oligonucleotides (oligos) and primers for Triplex-Seq

The developed Triplex-Seq assay requires oligonucleotides (oligos) which are referred to as triplex-forming oligos (TFOs), triplex target sites (TTS), DNA adapters which are ligated to TFOs in the downstream protocol and primers for PCR amplification. In the next sections, detailed descriptions of the design of all oligos, adapters and primers are given.

#### Design of triplex forming oligonucleotides (TFOs)

The triplex-forming oligos (TFOs) were designed with the following features and were ordered as single-stranded, desalted, non-modified DNA (ssDNA) oligos from integrated DNA technologies (IDT). Each TFO consists of two parts: (i) a common capture sequence and (ii) a triplex-forming sequence:

1. The 19 nt long capture sequence consists of fixed bases and serves as a platform for PCR amplification to enrich the TFOs. The capture sequence is the same for all TFOs that were tested in this study.
2. For the positive controls for triplex formation, the 20-30 nt long triplex-forming sequence consists of either adenine and guanine bases (anti-parallel TFO)^1,2^ or cytosine and thymine bases (parallel TFO)^3^. The TFO libraries were synthesized using the mixed bases tool (standard) from IDT. Mixed base oligos are synthesized according to the International Union of Pure and Applied Chemistry (IUPAC) convention. During chemical synthesis, each mixed base is integrated with percentages between 25 % (for ‘N’ mixed bases), 33 % (for ‘D’ and ‘B’ mixed bases) and 50 % (for ‘R’, ‘S’, ‘Y’, ‘W’, ‘M’, ‘K’ mixed bases) for every choice of base. Each TFO contains 6 fixed bases (with exception of the N-TFO library which contains 7 fixed bases) and 14 mixed bases (with exception of the N-TFO library which contains 13 mixed bases). The 6-7 fixed bases provide a TFO-library barcode, for the case that two or more libraries are sequenced in the same sequencing run.

In Table 1-3, a list of all TFOs (Table 1), TFO libraries (Table 2) and verification TFOs (Table 3) are shown. Each table includes the TFO name, sequence and number of variants per TFO library. In Table 1, the TFOs that served as positive controls are shown. Here, they were used to verify triplex formation and were used in the *in vitro* Triplex-Seq protocol to confirm that the conditions are optimal to induce triplex formation. All TFOs that were used to confirm triplex formation *in vitro* were either adapted from literature or designed as part of this study with the support of the prediction software Triplexator^4^.

| **Table 1\|** **Literature TFOs and control TFOs for *in vitro* Triplex-Seq**. | | | |
| --- | --- | --- | --- |
| **name** | **TFO sequence (5' –> 3')** | **length [nt]** | **reference** |
| TFO_AG30 | AGGAAGGGGGGGGTGGTGGGGGAGGGGGAG | 30 | ^1,2^ |
| TFO_TC | CCTCTTCCTCCTCTTCCTTTCTC | 23 | ^3^ |
| TFO_p_G20 | CTTCAGCTTGGCGGTCTGGCTTTTCTCTTTTCTTCTTTT | 39 | this work |
| TFO_ap_G80 | CTTCAGCTTGGCGGTCTGGGGAGGGGGAGGGGGGGGAGG | 39 | this work |

The TFO libraries were constructed as described above and are listed in Table 2 including the fixed bases, the mixed bases used, the sequence as it was ordered, the total length of the TFO and the number of variants for each library. Each TFO library was synthesized according to the IUPAC convention. The name and the description indicate the nature of the TFO (e.g. which mixed base was used, how many fixed bases can be found within the TFO and whether a capture sequence was attached), the sequence, length and the number of variants for each library follow the description.

| **Table 2\| List of TFO libraries for the Triplex-Seq approaches**. | | | |
| --- | --- | --- | --- |
| **name** | **TFO sequence (5' –> 3')** | **length [nt]** | **variants** |
| oTFO_3D | CTTCAGCTTGGCGGTCTGGCATCCTGADDDCTGACATGC | 39 | 27 |
| oTFO_3B | CTTCAGCTTGGCGGTCTGGCATCGTAABBBAAGACATGC | 39 | 27 |
| oTFO_5D | CTTCAGCTTGGCGGTCTGGCATCCTGDDDDDTGACATGC | 39 | 243 |
| oTFO_5B | CTTCAGCTTGGCGGTCTGGCATCGTGBBBBBAGACATGC | 39 | 243 |
| oTFO_M | CTTCAGCTTGGCGGTCTGGTMAMMGMMMMMCTMMMMAMM | 39 | 1.6E+04 |
| oTFO_R | CTTCAGCTTGGCGGTCTGGRRGRTRRRARRRRRGRRART | 39 | 1.6E+04 |
| oTFO_K | CTTCAGCTTGGCGGTCTGGCKGKKTKKKKKGTKKKKAKK | 39 | 1.6E+04 |
| oTFO_N | CTTCAGCTTGGCGGTCTGGCNANNTNNNNNTGNNNNCNG | 39 | 2.7E+08 |
| oTFO_7D | CTTCAGCTTGGCGGTCTGGCATCCTDDDDDDDGACATGC | 39 | 2187 |
| oTFO_7B | CTTCAGCTTGGCGGTCTGGCATCGTBBBBBBBGACATGC | 39 | 2187 |
| oTFO_9D | CTTCAGCTTGGCGGTCTGGCATCCDDDDDDDDDACATGC | 39 | 1.97E+04 |
| oTFO_9B | CTTCAGCTTGGCGGTCTGGCATCGBBBBBBBBBACATGC | 39 | 1.97E+04 |
| oTFO_W | CTTCAGCTTGGCGGTCTGGTWAWWWWWTWWWWWAWWTWA | 39 | 1.6E+04 |

Furthermore, single-variant TFOs were ordered based on the enrichment of sequences after the Triplex-Seq analysis. In Table 3 below, all ordered and tested TFO sequences are listed. Following NGS analysis of the Triplex-Seq reads that were enriched in the downstream Triplex-Seq protocol, several single-variants of the most enriched hits (“positive TFOs”) and least enriched hits (“negative TFOs”) of the triplex band from the *in vitro* Triplex-Seq protocol were ordered and tested. The names in the first column describe from which (i) TFO library and (i) condition the sequence of the TFO variant is derived from. The description in the fourth columns indicates in which figure in the study the results are shown.

| **Table 3\| List of TFOs used in verification experiments of Triplex-Seq**. | | | |
| --- | --- | --- | --- |
| **name** | **TFO sequence (5' –> 3')** | **length [nt]** | **description** |
| M_pH5_triple | AACGACCCCCGACCCACCCCC | 21 | Extended Data Figure 1, #1 |
| M_pH7_triple | AACACCCCCACCCCAACC | 18 | Extended Data Figure 1, #2 |
| M_hit2_pH7 | TAAAAGAAAAACTACACACC | 20 | Extended Data Figure 1, #3 |
| M_hit1_pH5 | TAAAAGAAAAACTACAAACA | 20 | Extended Data Figure 1, #4 |
| M_hit2_pH5 | TCAAAGCAACCCTCCACAAA | 20 | Extended Data Figure 1, #5 |
| N_pos3_pH7 | CTAGTTGGGGGTGGGGGCGG | 20 | Extended Data Figure 2, enriched |
| N_neg1_pH7 | CGAGGTTATGATGAAACCGG | 20 | Extended Data Figure 2, non-enriched |
| DRIMust_pH5 | GTGGGAGCGG | 10 | Figure 4, #1 |
| pH5_75%_double | GTGGGGGCGTGGGGGT | 16 | Figure 4, #2 |
| DRIMust_G_pH7 | GGGGTGGGGGC | 11 | Figure 4, #3 |
| DRIMust_A_pH7 | GGGGTGGGAGC | 11 | Figure 4, #4 |
| DRIMust_2 _pH7 | GGGGTGGGGGCGGGGTGGGGACG | 23 | Figure 4, #5 |

#### Design of triplex target sites (TTS)

The triplex-target sites (TTS) were designed based on sequences found in literature and were tested *in vitro*. We initially used the original TTS sequences as they have been described in the publications. After verification that they worked we chose to continue with two TTSs in the Triplex-Seq process. For the purpose of the Triplex-Seq protocol, we expanded the sequence of the original TTS by approx. 20 nt on each side (5' and 3'). The TTSs were generated by annealing single-stranded oligos (95 ºC for 2 minutes, cool-down to RT over a course of 45 minutes) and the sequence of each oligo is shown in Table 4. In addition to the TTSs that were based on literature sequences (thus termed positive controls), we also designed new TTSs with increasing frequency of guanines within the sequence, starting from 20 % guanines up to 80 % guanines. The design of these TTSs as well as corresponding TFOs were supported by the Triplexator software. Use of the software is described in the bioinformatics section.

| **Figure 4\| List of triplex-target sites used for the *in vitro* Triplex-Seq setup.** | | | |
| --- | --- | --- | --- |
| **name** | **TFO sequence (5' –> 3')** | **length [nt]** | **used in Fig.** |
| oTTS-G20fw | GGCCGCTTTTCTTTTCTCTTTTCTTCTTTTTTCTTTGACGT | 41 | 4 |
| oTTS-G20rev | CAAAGAAAAAAGAAGAAAAGAGAAAAGAAAAGC | 33 | 4 |
| oTTS-G33fw | GGCCGCTCTTCTTTTCTTCTTTCTTCCTTCTTCCTTGACGT | 41 | 4 |
| oTTS-G33rev | CAAGGAAGAAGGAAGAAAGAAGAAAAGAAGAGC | 33 | 4 |
| oTTS-G53fw | GGCCGCTCCTCCTCCCTTCTTTCTTCCTTCTTCCCTGACGT | 41 | 4 |
| oTTS-G53rev | CAGGGAAGAAGGAAGAAAGAAGGGAGGAGGAGC | 33 | 4 |
| oTTS-G75fw | GGCCGCTCCCCCTCCCTCCTTTCCTCCTCCCCCCCTGACGT | 41 | 4 |
| oTTS-G75rev | CAGGGGGGGAGGAGGAAAGGAGGGAGGGGGAGC | 33 | 4 |
| oTTS-G84fw | GGCCGCTCCCCCTCCCCCCCCTCCCCCTCCCCCCCTGACGT | 41 | 4 |
| oTTS-G84rev | CAGGGGGGGAGGGGGAGGGGGGGGAGGGGGAGC | 33 | 4 |
| oTTS-2_80_fw | GTATCGTAATACGATGCGCATGCTACGTTGGAGAAGGAGGAGAAGGAAAGAGTCCTCTATACGCAGACTCAAGCTGACC | 79 | 2 + 3 |
| oTTS-2_80_rev | GGTCAGCTTGAGTCTGCGTATAGAGGACTCTTTCCTTCTCCTCCTTCTCCAACGTAGCATGCGCATCGTATTACGATAC | 79 | 2 + 3 |
| oTTS-1_80_fw | GTATCGTAATACGATGCGGTTCGAATCCTTCCCCCCCCACCACCCCCTCCCCCTCCAGACTCAAGCTGACC | 71 | 2 + 3 |
| oTTS-1_80_rev | GGTCAGCTTGAGTCTGGAGGGGGAGGGGGTGGTGGGGGGGGAAGGATTCGAACCGCATCGTATTACGATAC | 71 | 2 + 3 |
| TTS-1_37 (fw) | GTTCGAATCCTTCCCCCCCCACCACCCCCTCCCCCTC | 37 | ED1 |
| TTS-1_37 (rev) | GAGGGGGAGGGGGTGGTGGGGGGGGAAGGATTCGAAC | 37 | ED1 |
| TTS-2_45 (fw) | CATGCTACGTTGGAGAAGGAGGAGAAGGAAAGAGTCCTCTATACG | 45 | ED1 |
| TTS-2_45 (rev) | CGTATAGAGGACTCTTTCCTTCTCCTCCTTCTCCAACGTAGCATG | 45 | ED1 |
| Forward and reverse sequences of TTS oligos are shown. ap, antiparallel (pH 7); p, parallel (pH 5); fw, forward; rev, reverse; ED, Extended Data | | | |

#### Design of primers and other oligos for sequencing

For the preparation of the TFO sequences for Illumina sequencing (see detailed experimental setup below), a single-stranded DNA (ssDNA) adapter was designed for ssDNA ligation as well as primers for PCR amplification of TFO sequences and simultaneous addition of Illumina sequences. A full list of primers and adapter sequences is shown in Table 5 and highlights the sequences as well as modifications of the primers. All primers were ordered as desalted ssDNA oligos from IDT. Deviations of standard primers are mentioned in description.

| **Table 5\| Primers and oligos for Triplex-Seq protocol**. | |
| --- | --- |
| **oligo name** | **sequence (5' –> 3')** |
| General Illumina sequence | CAAGCAGAAGACGGCATACGAGATNNNNNNGTGACTGGAGTTCAGACGTGTGCTC (N = #1-#35) |
| Illumina Index #1 | CGTGAT |
| Illumina Index #2 | ACATCG |
| Illumina Index #3 | GCCTAA |
| Illumina Index #4 | TGGTCA |
| Illumina Index #5 | CACTGT |
| Illumina Index #6 | ATTGGC |
| Illumina Index #7 | GATCTG |
| Illumina Index #8 | TCAAGT |
| Illumina Index #9 | CTGATC |
| Illumina Index #10 | AAGCTA |
| Illumina Index #11 | GTAGCC |
| Illumina Index #12 | TACAAG |
| Illumina Index #13 | TTGACT |
| Illumina Index #14 | GGAACT |
| Illumina Index #15 | TGACAT |
| Illumina Index #16 | GGACGG |
| Illumina Index #17 | CTCTAC |
| Illumina Index #18 | GCGGAC |
| Illumina Index #19 | TTTCAC |
| Illumina Index #20 | GGCCAC |
| Illumina Index #21 | CGAAAC |
| Illumina Index #22 | CGTACG |
| Illumina Index #23 | CCACTC |
| Illumina Index #24 | GCTACC |
| Illumina Index #25 | ATCAGT |
| Illumina Index #26 | GCTCAT |
| Illumina Index #27 | AGGAAT |
| Illumina Index #28 | CTTTTG |
| Illumina Index #29 | TAGTTG |
| Illumina Index #30 | CCGGTG |
| Illumina Index #31 | ATCGTG |
| Illumina Index #32 | TGAGTG |
| Illumina Index #33 | CGCCTG |
| Illumina Index #34 | GCCATG |
| Illumina Index #35 | AAAATG |
| ssDNA adapter | /5Phos/AGATCGGAAGAGCACACGTCTGAACTCCAGTCAC/3SpC3/ |
| Biot-TriSeqNGS-ddC | /5BiosG/GAAGTCGAACCGCCAGACC/3ddC/ |
| PE_forward | AATGATACGGCGACCACCGAGATCTACACTCTTTCCCTACACGACGCTCTTCCGATCTCTTTCCCTACACGACGCTCTTCCGATCTCTTCA |
